# Autophagy cargo receptors are secreted via extracellular vesicles and particles in response to endolysosomal inhibition or impaired autophagosome maturation

**DOI:** 10.1101/2021.08.12.456045

**Authors:** Tina A. Solvik, Tan A. Nguyen, Yu-Hsiu Tony Lin, Timothy Marsh, Eric J. Huang, Arun P. Wiita, Jayanta Debnath, Andrew M. Leidal

## Abstract

The endosome-lysosome (endolysosome) system plays central roles in both autophagic degradation and secretory pathways, including the exocytic release of extracellular vesicles and particles (EVPs). Although previous work has revealed important interconnections between autophagy and EVP-mediated secretion, our molecular understanding of these secretory events during endolysosome inhibition remains incomplete. Here, we delineate a secretory autophagy pathway upregulated in response to endolysosomal inhibition that mediates the EVP-associated extracellular release of autophagic cargo receptors, including p62/SQSTM1. This extracellular secretion is highly regulated and critically dependent on multiple ATGs required for the progressive steps of early autophagosome formation as well as Rab27a-dependent exocytosis. Furthermore, the disruption of autophagosome maturation, either due to genetic inhibition of the autophagosome-to-autolyosome fusion machinery or blockade via the SARS-CoV2 viral protein ORF3a, is sufficient to induce robust EVP-associated secretion of autophagy cargo receptors. Finally, we demonstrate that this ATG-dependent, EVP-mediated secretion pathway buffers against the intracellular accumulation of autophagy cargo receptors when classical autophagic degradation is impaired. Based on these results, we propose that secretory autophagy via EVPs functions as an alternate route to clear sequestered material and maintain proteostasis in response to endolysosomal dysfunction or impaired autophagosome maturation.

## Introduction

Although autophagy is classically viewed as a lysosomal degradation pathway, increasing evidence implicates autophagy-related genes (ATGs) in the secretion of inflammatory molecules (Dupont et al., 2011; Lock et al., 2014), tissue repair factors (DeSelm et al., 2011) and bactericidal enzymes (Bel et al., 2017), and extracellular vesicles (EVs) (Guo et al., 2017; Leidal et al., 2020; Murrow et al., 2015). Recently, we described an autophagy-related secretory pathway called LC3-dependent EV loading and secretion (LDELS), which captures proteins at late endosomes and facilitates their secretion outside the cell (Leidal et al., 2020). These expanding roles for autophagy pathway components in secretion, collectively termed secretory autophagy, poignantly suggest that cellular catabolism and secretion are functionally coordinated within cells. Nevertheless, the mechanisms orchestrating secretory autophagy versus classical degradative autophagy remain poorly understood as well as how this interplay contributes to proteostasis and cellular quality control.

Notably, recent studies demonstrate that endolysosome impairment profoundly impacts the dynamics of both autophagy-dependent degradation and EV secretion. Lysosomal inhibition via Bafilomycin A1 and chloroquine treatment impairs autophagy and stimulates EV secretion (Cashikar and Hanson, 2019; Mauthe et al., 2018; Ortega et al., 2019). Moreover, chemical inhibitors targeting PIKfyve and VPS34, two PI3K kinases important for endosomal trafficking and autophagy, disrupt autophagy-dependent protein turnover and promote secretion of various autophagy components in EV fractions (Hessvik et al., 2016; Miranda et al., 2018). These observations suggest that the status of the endolysosomal system influences secretory autophagy via EV release, which bears fundamental significance for understanding various pathological conditions and therapies that impair endolysosome function. Importantly, many pathogens specifically impede endolysosome function to prevent clearance by the autophagy pathway. For example, lysosomal disruption during uropathogenic *E. coli* infection triggers autophagy-dependent expulsion of bacteria in EVs, whereas many RNA viruses exploit autophagic membranes that accrue in response to impaired lysosomal degradation for viral replication and exocytosis (Miao et al., 2015; Teo et al., 2021). Endolysosome deregulation is also common in age-related diseases including neurodegeneration and autophagy-dependent secretion in diseased neurons may contribute to the spread of aggregation-prone proteins (Ejlerskov et al., 2013; Minakaki et al., 2018; Nilsson et al., 2013). Finally, the lysosomal inhibitor hydroxychloroquine (HCQ) has been extensively employed in multiple clinical oncology trials in order to target the heightened dependence of cancer cells on the autophagy-lysosome pathway for growth and survival (Amaravadi et al., 2019). Nevertheless, whether and how endolysosome impairment in these contexts contributes to secretory autophagy, and more specifically to EV-associated secretion, remains largely unknown.

Here, we demonstrate that lysosomal inhibition robustly induces ATG-dependent secretion of autophagy components and autophagy cargo receptors in EV fractions both *in vitro* and *in vivo*. Remarkably, autophagy cargo receptors secreted in EV-associated fractions during lysosome inhibition are largely unprotected from proteolytic cleavage, suggesting that in contrast to the LDELS pathway, these proteins are not selectively loaded into EVs. Rather, they are released outside the cell in tight association with EVs but as part of a broader fraction of nanoparticles, termed collectively as extracellular vesicles and particles (EVPs) (Zhang et al., 2018). This extracellular secretion is highly regulated and critically dependent on multiple ATGs required for the progressive steps in autophagosome formation. Furthermore, disruption of autophagosome maturation, due to genetic loss of the autophagosome-to-autolyosome fusion machinery or blockade via the SARS-CoV2 viral protein ORF3a, is sufficient to induce robust EVP-associated secretion of autophagy cargo receptors. Finally, the suppression of this secretory pathway promotes intracellular accumulation of individual cargo receptors suggesting that when autophagy-dependent degradation is impaired due to endolysosome impairment, the autophagy machinery diverts autophagic cargo receptors for secretion outside the cell via EVPs.

## Results

### Inhibition of lysosome acidification promotes the EVP-associated release of core autophagy machinery and degradative cargo

To ascertain how lysosome impairment affects ATG-dependent EV secretion, we treated wild-type and ATG7-deficient HEK293T cells with the vacuolar-type ATPase (v-ATPase) inhibitor Bafilomycin A1 (BafA1), which disrupts lysosome acidification, and evaluated its impact on protein cargo released in small EV fractions (Fig. 1A). Treatment of serum starved wild-type cells with BafA1 resulted in robust increases in the secretion of endosomal proteins in EV fractions including the Rab GTPases Rab5a and Rab7a, markers of early and late endosomes, respectively (Fig. 1B). In contrast, endosomal protein secretion was largely unaltered in BafA1-treated, ATG7-deficient cells, which are defective for the lipid conjugation of MAP1LC3B (LC3) family proteins that is essential for both classical autophagy and emerging secretory autophagy pathways, including LC3-Dependent EV Loading and Secretion (LDELS) (Nieto-Torres et al., 2021). These results broached that the autophagy machinery facilitates the diversion of endocytic intermediates for extracellular secretion during lysosome inhibition. Importantly, Rab5a and Rab7a secretion during BafA1 treatment correlated with impaired autophagosome maturation, supported by BafA1-induced accumulation of yellow puncta (mCherry-positive; GFP-positive), which delineate early, immature autophagosomes in cells expressing the mCherry-GFP-LC3B autophagic flux reporter (Sup. Fig. 1A,B) (Pankiv et al., 2007). We also observed robust increases in the co-localization of LC3 and the late endosome marker CD63 in response to BafA1 (Sup. Fig. 1C, D), suggesting that this autophagy protein was redirected towards late endosomes when lysosome acidification was impaired. Although nanoparticle tracking analysis of conditioned media revealed that BafA1 did not significantly impact the overall levels and mean particle size of secreted EVPs, we did note increases in specific nanoparticle populations measuring 110 nm, 125 nm and 140 nm diameter (Sup. Fig. 1E,F).

**Figure 1.**
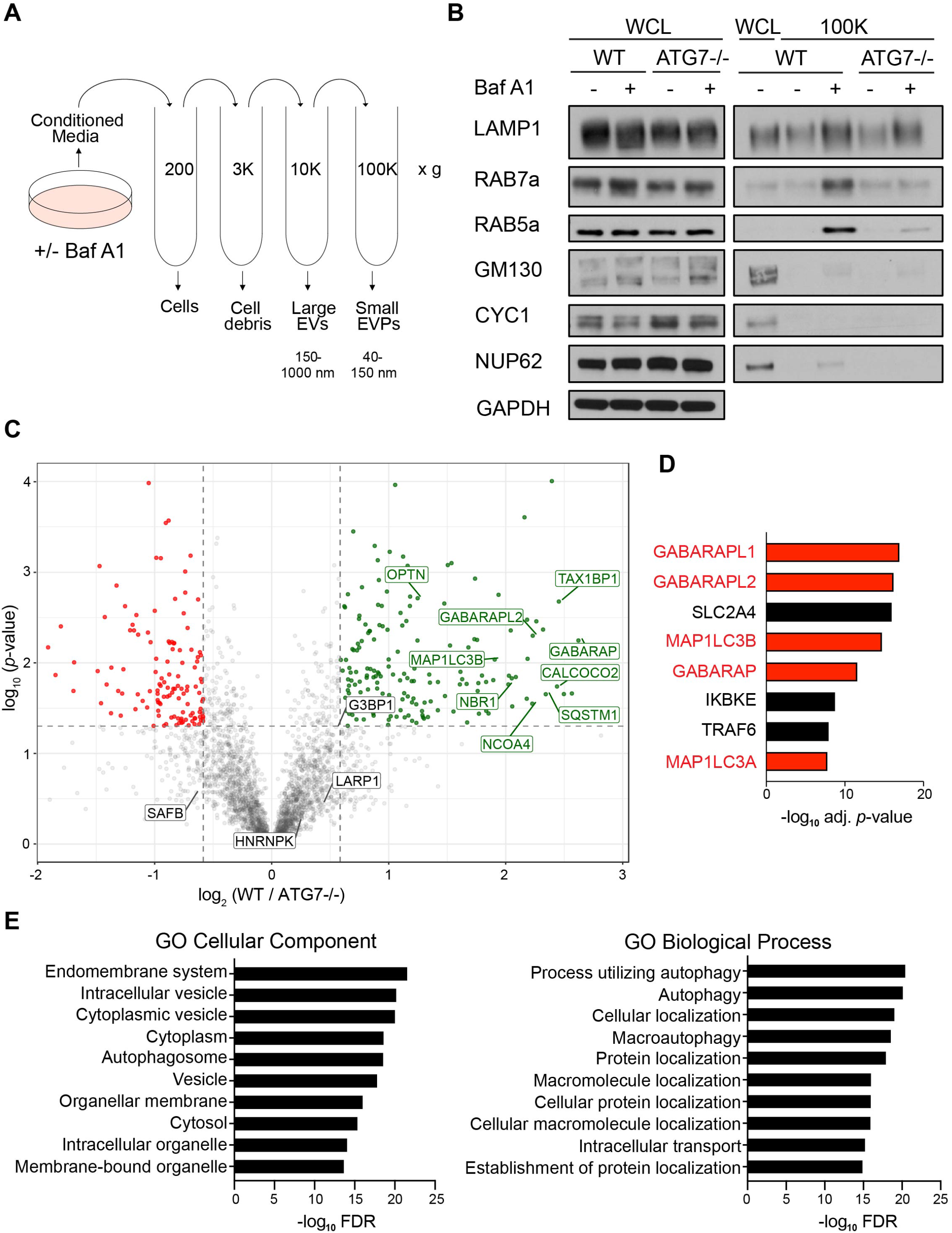
BafA1-induced disruption of lysosome acidification promotes ATG-dependent secretion of autophagy cargo receptors via EVPs. **(A)** Schematic detailing the differential centrifugation protocol employed to isolate small EVP-enriched fractions from cells treated with BafA1 or vehicle. **(B)** Whole cell lysate (WCL; left) and 100,000g EVP fractions (100K; right) from wild-type (WT) and ATG7-/- serum starved HEK293T cells treated with 20 nM BafA1 or vehicle were collected, normalized to cell number and immunoblotted to detect the indicated proteins (n = 3). **(C)** A volcano plot of the proteins identified in 100,000g EVP-enriched fractions from BafA1 treated WT and ATG7−/− HEK293T cells quantified by TMT mass spectrometry. TMT-labelled proteins are plotted according to their −log_10_ P values as determined by two-tailed t-test and log^2^ fold enrichment (WT/ATG7−/−; n = 3). Grey dots: proteins not relatively enriched in EVPs from BafA1 treated WT or ATG7−/− cells identified with P > 0.05 and/or log_2_ fold change between −0.5 and 0.5. Green dots: proteins significantly enriched in EVPs from BafA1 treated WT cells relative to treated ATG7−/− cells. Red dots: proteins significantly enriched in EVPs from BafA1 treated ATG7−/− cells relative to treated WT cells. **(D)** A ranked list of the proteins with the greatest connectivity to the 182 proteins enriched in EVPs from BafA1 treated WT cells relative to treated ATG7−/− cells. Statistical significance was calculated in Enrichr by a one-way Fisher’s exact test. LC3 family members are highlighted in red. PPI, protein–protein interaction. **(E)** Gene Ontology (GO) enrichment analysis of the 182 proteins enriched in EVPs from BafA1 treated WT cells relative to treated ATG7−/− cells with the top terms for cellular component (left) and biological processes (right) plotted according to −log_10_ False Discovery Rate (FDR). Statistical significance was calculated by a one-way Fisher’s exact test.

To further define the ATG-dependent secreted proteome during lysosome inhibition in an unbiased manner, we performed Tandem Mass Tag (TMT)-based quantitative proteomics comparing small EV fractions purified via differential centrifugation (100,000g) from the conditioned media of wild-type and ATG7-deficient cells treated with BafA1. Importantly, while these 100,000g fractions are enriched for small EVs, they contain a broader array of nanoparticles and proteins, collectively termed extracellular vesicles and particles (EVPs) (Zhang et al., 2018). Overall, we identified 182 proteins enriched in the EVP fractions from BafA1-treated wild-type versus ATG7-deficient cells (Fig. 1C). Many were core autophagy machinery proteins including LC3/ATG8 family members as well as components of complexes that regulate discrete steps of autophagosome formation. Importantly, we also identified numerous autophagy cargo receptors, proteins critical for selectively capturing cargo for autophagic degradation as well as serving as scaffolds for cellular signaling (Johansen and Lamark, 2020). These included p62/SQSTM1, NBR1, OPTN and NDP52 (Fig. 1C). Notably, five LC3/ATG8 family members ranked amongst the top proteins connected to the 182 proteins enriched in the BafA1-induced ATG7-dependent EV proteome (Fig. 1D). Moreover, gene ontology (GO) analyses highlighted a profound enrichment in proteins that localize to the endomembrane system and proteins with roles in autophagy (Fig. 1E, F). Interestingly, the ATG7-dependent EVP proteome from BafA1 treated cells was quite distinct from the recently defined LDELS proteome; there was no evidence for functional enrichment of RBPs, including SAFB, HNRNPK, LARP1 and G3BP1, in control versus ATG7 deficient cells treated with Baf A1 (Fig. 1C). Overall, the BafA1-induced ATG7-dependent EVP proteome shared twelve proteins in common with the BirA*-LC3B-labelled secretome, fifty four proteins with the ATG7 and ATG12-dependent EV proteome previously described for LDELS, and two proteins with the ATG5-dependent secretome from murine macrophages (Fig. 2A-C) (Kimura et al., 2017; Leidal et al., 2020). Remarkably, the ATG7-dependent EVP proteome secreted by BafA1-treated cells shared numerous common proteins with proteomic datasets derived from studies of degradative autophagy (Fig. 2D, E) (Behrends et al., 2010; Mancias et al., 2014). Based on these results, we hypothesized that lysosome impairment diverts proteins residing in autophagosomes, but originally destined for degradation, for secretion outside the cell via EVPs.

**Figure 2.**
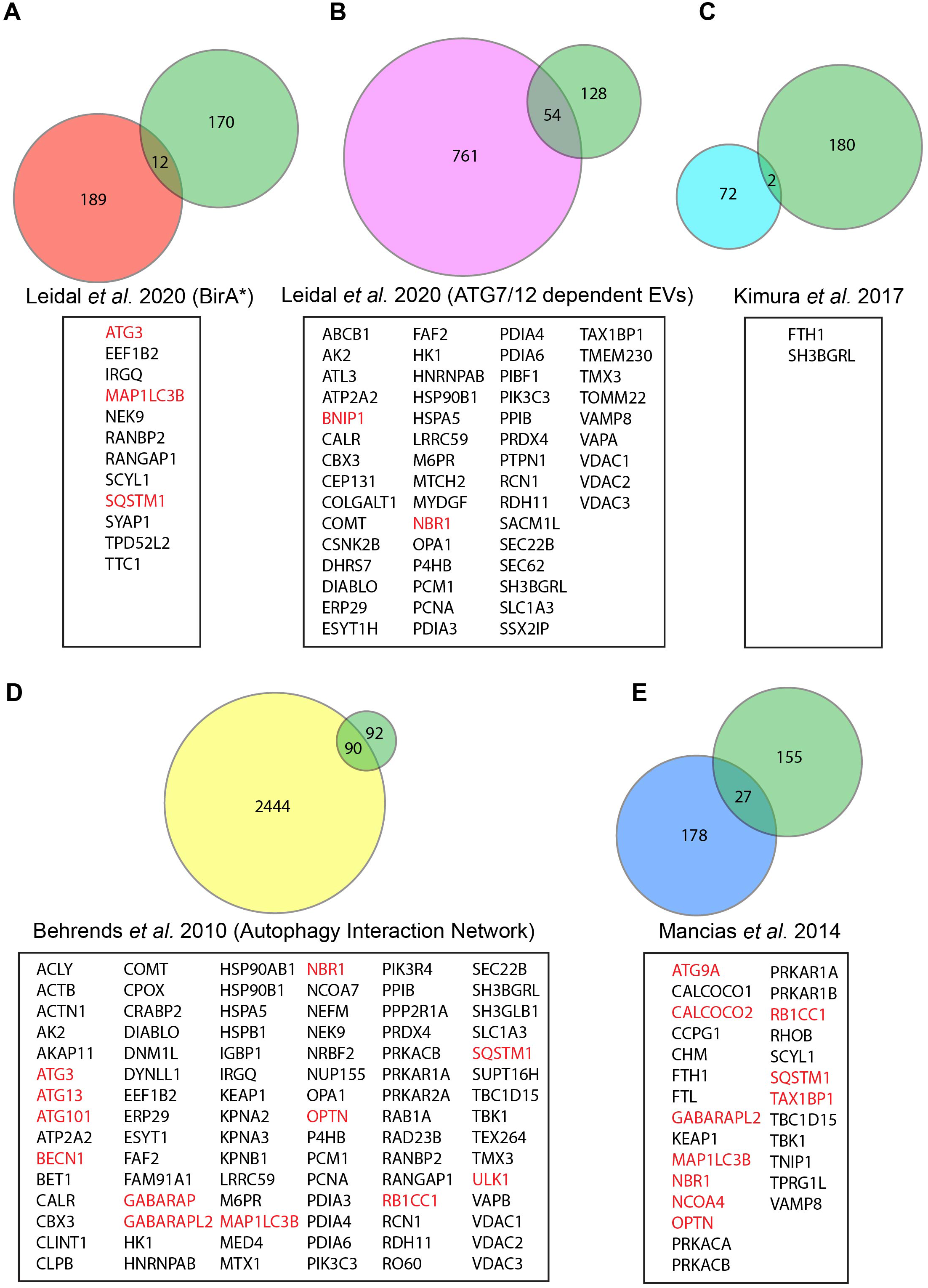
The ATG-dependent EVP secretome from BafA1 treated cells overlaps with previous proteomic analyses of the autophagy pathway. **(A)** Venn diagram showing the overlap of ATG-dependent EVP secretion candidates from BafA1 treated cells with the class I and class II BirA*-LC3B labelled secretome defined in Leidal *et al*. 2020. **(B)** Venn diagram showing the overlap of ATG-dependent EVP secretion candidates from BafA1 treated cells with the ATG7 and ATG12-dependent EV secretome defined in Leidal *et al*. 2020. **(C)** Venn diagram showing the overlap of ATG-dependent EVP secretion candidates from BafA1 treated cells with the ATG7 and ATG12-dependent EV secretome defined in Leidal *et al*. 2020. **(D)** Venn diagram showing the overlap of ATG-dependent EVP secretion candidates from BafA1 treated cells with the ATG5-dependent bone-marrow derived macrophage secretome defined in Kimura *et al*. 2017. **(E)** Venn diagram showing the overlap of ATG-dependent EVP secretion candidates from BafA1 treated cells with the complete Autophagy Interaction Network (AIN) defined in Behrends *et al*. 2010. **(F)** Venn diagram showing the overlap of ATG-dependent EVP secretion candidates from BafA1 treated cells with the autophagosome enriched proteome defined in Mancias *et al*. 2014. Core autophagy machinery and autophagy cargo receptors are highlighted in red.

### Lysosome inhibition promotes EVP secretion of autophagy cargo receptors *in vitro* and *in vivo*

We next assessed whether specific ATG7-dependent secretion candidates identified via TMT-MS were release in association with EVPs in response to lysosome inhibition. Small EVP fractions from serum starved wild-type cells treated in the absence and presence of BafA1 were isolated via differential centrifugation and probed for autophagy pathway components and markers of EVs. Importantly, the 100,000g fraction from BafA1 treated cells was highly enriched in lipidated LC3 (LC3-II) and multiple autophagy cargo receptors including p62/SQSTM1, NBR1, OPTN and NDP52. In contrast, BafA1 treatment had a negligible impact on the secretion of TSG101, ALIX and CD9, classical protein markers of EV populations (Fig. 3A, B). To test whether this effect was a more general response to lysosome inhibition and impaired autophagosome maturation, we assessed EV secretion in cells treated with chloroquine, a lysosomotropic anti-malarial agent used to target the autophagy-lysosome pathway for cancer therapy (Amaravadi et al., 2019). Chloroquine (CQ), similar to BafA1, robustly increased extracellular secretion of LC3-II and autophagy cargo receptors in EV fractions isolated from conditioned media without significantly altering the secretion of EV marker proteins (Fig. 3C, D). Importantly, enhanced secretion of LC3-II and autophagy cargo receptors during lysosome inhibition did not correlate with increased cell death and was observed in a diverse array of BafA1 treated cells, including murine and human breast cancer cell lines (Sup. Fig. 2 and Sup. Fig. 3). Finally, upon treatment of GFP-LC3 transgenic mice with CQ and purification of EVP fractions from the blood plasma of these animals, we observed increased in vivo extracellular secretion of both LC3 and the cargo receptors p62/SQSTM1 and NBR1 in CQ-treated mice compared to untreated controls (Fig. 3E,F). Overall, these findings confirm that pharmacological inhibition of the lysosome results in the increase extracellular secretion of LC3 and autophagy cargo receptors both *in vitro* and *in vivo*.

**Figure 3.**
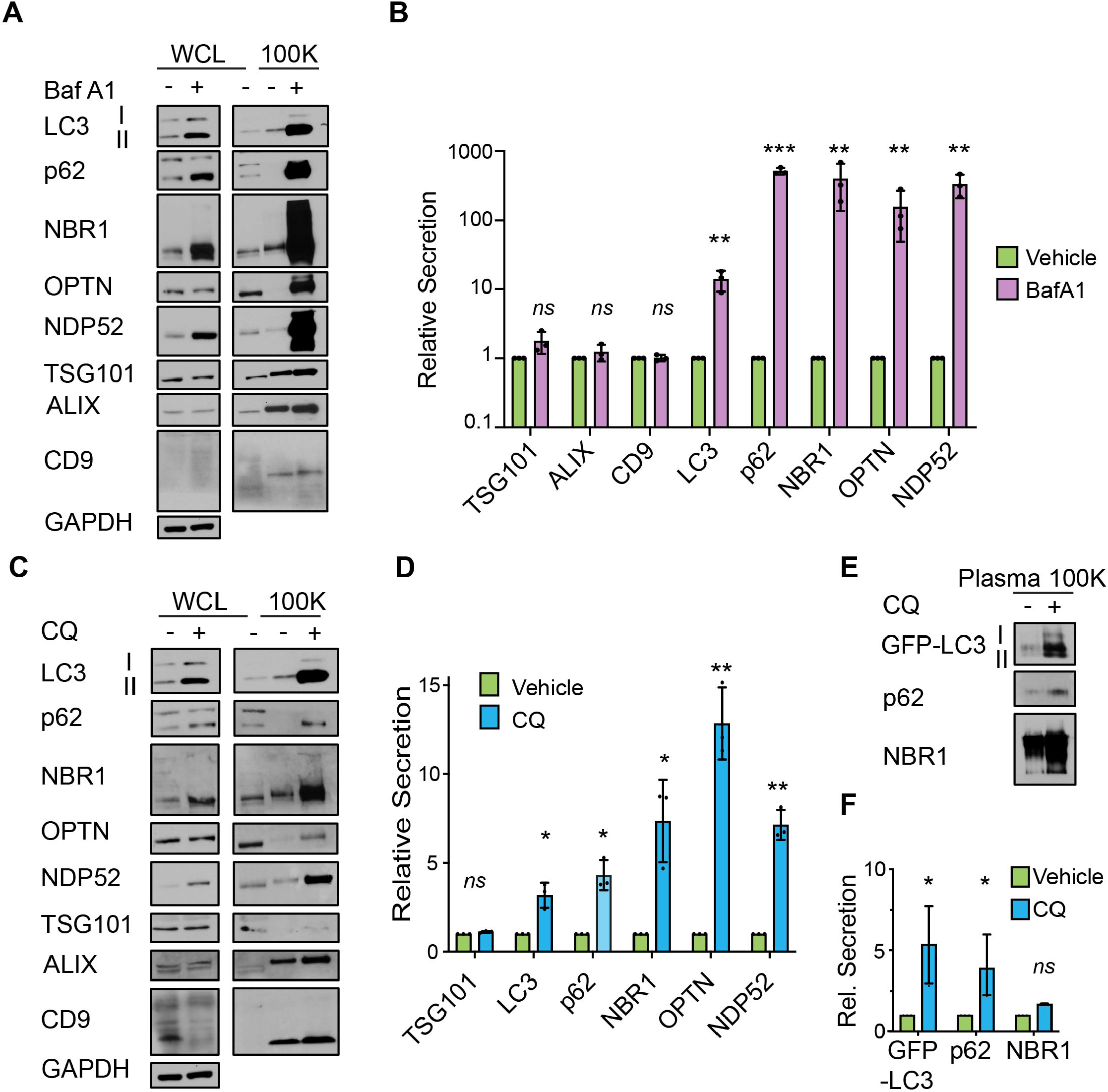
Lysosomal inhibition broadly promotes secretion of autophagy cargo receptors via EVPs. **(A)** Whole cell lysate (WCL; left) and 100,000g EVP fractions (100K; right) from serum starved HEK293Ts treated with vehicle or 20 nM BafA1 for 16h were collected and immunoblotted to detect the indicated proteins (n = 3). **(B)** Quantification of the indicated proteins in EVP fractions from BafA1 treated cells relative to vehicle controls in Panel A. (mean ± s.e.m.; n = 3; ns=not significant, p<0.01= **, p<0.005=***). Statistical significance was calculated by one-way ANOVA coupled with Tukey’s post hoc test. **(C)** WCL (left) and 100K fractions (right) from serum starved HEK293Ts treated with vehicle or 25 µM chloroquine (CQ) for 16 h were collected and immunoblotted to detect the indicated proteins (n = 3). **(D)** Quantification of the proteins in EVP fractions from CQ treated cells relative to untreated controls in Panel C (mean ± s.e.m.; n = 3; ns=not significant, p<0.05= *, p<0.01=**). Statistical significance was calculated by one-way ANOVA coupled with Tukey’s post hoc test. **(E)** Plasma EVPs from mice treated with vehicle or 60 mg/kg chloroquine (CQ) for 3 consecutive days were collected and immunoblotted for the levels of GFP-LC3, p62 and NBR1. **(F)** Quantification of the indicated proteins in plasma EVPs from mice treated with vehicle or CQ from Panel E (mean ± s.e.m.; n = 3; ns=not significant, p<0.05= *). Statistical significance was calculated by one-way ANOVA coupled with Tukey’s post hoc test.

### Autophagy cargo receptors are not extracellularly secreted within protease-protected vesicular intermediates

Our group recently revealed a novel role for the autophagy conjugation machinery in specifying RNA-binding proteins for secretion within small EVs (Leidal et al., 2020). To more precisely determine whether LC3 and autophagy cargo receptors are specifically released via small EVs during lysosome impairment, we performed serial differential centrifugation of conditioned media from serum starved, BafA1 treated cells to isolate fractions enriched in large EVs (10,000g), small EVs and particles (EVPs, 100,000g) and soluble proteins, which were precipitated from the remaining sample. Notably, the 100,000g fraction from BafA1 treated cells was highly enriched in LC3, p62, NBR1, OPTN and NDP52, whereas the other fractions contained significantly less LC3 and near undetectable levels of autophagy cargo receptors (Fig. 4A, B). Upon further purifying EVs from BafA1 treated cells via linear sucrose density gradient ultracentrifugation, we found that LC3-II and autophagy cargo receptors in these isolates co-fractionated with the EV-associated tetraspanin CD9 (Fig. 4C, D). Finally, we performed protease protection of 100,000g fractions to assess whether autophagy cargo receptors released during lysosome impairment were localized within the lumens of small EVs. However, in contrast to the EV luminal marker TSG101, the autophagy cargo receptors p62, NBR1 and OPTN within EV fractions were almost completely digested by trypsin in the absence of detergent (Fig. 4E, F). Together, these results indicated that autophagy cargo receptors released during lysosome inhibition are predominately associated with either the exterior surface of small EVs or other secreted particles (EVPs), rather than being directly packaged into the lumen of EVs.

**Figure 4.**
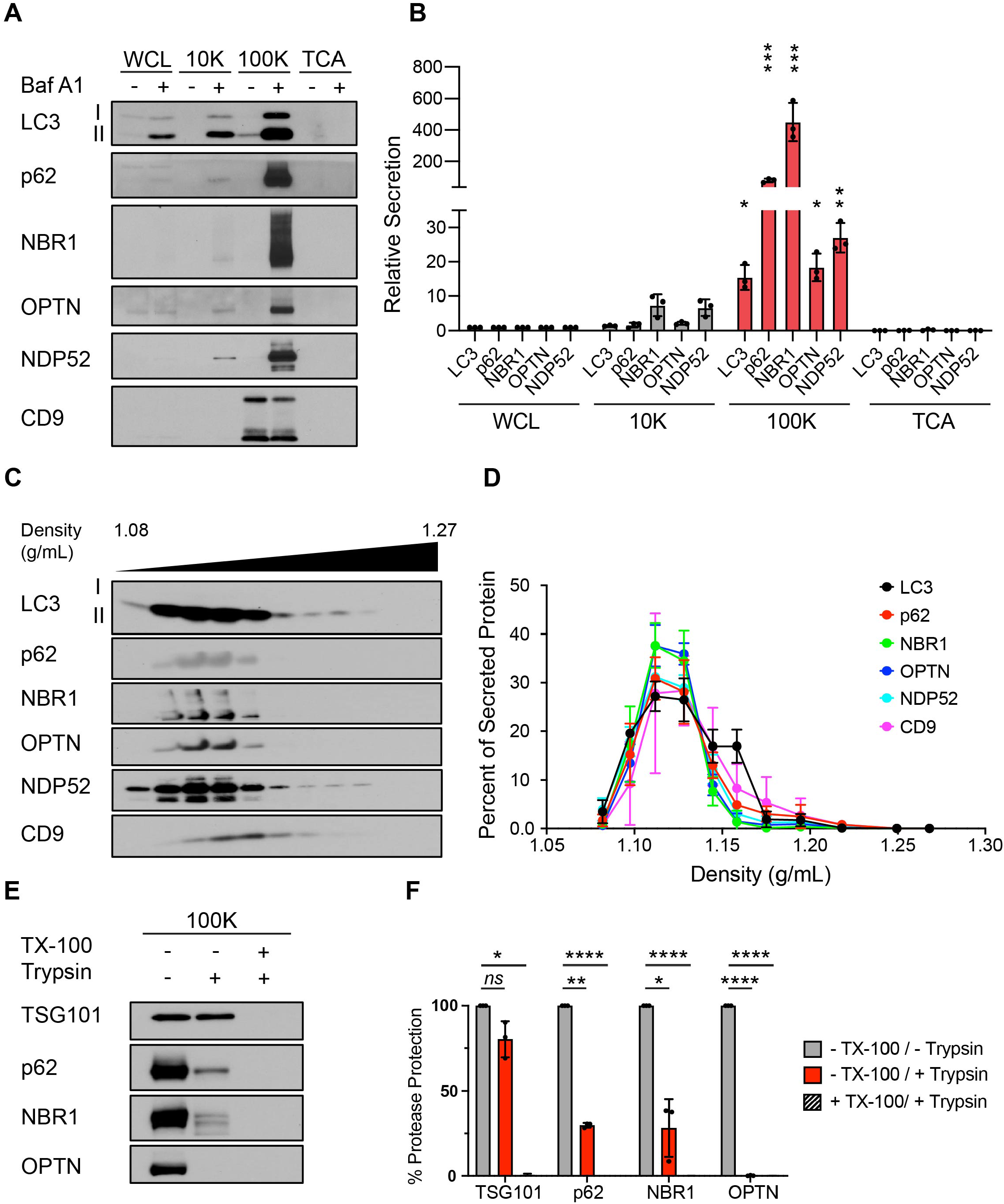
Autophagy cargo receptors are secreted as EVP-associated proteins in response to lysosome inhibition. **(A)** Whole cell lysate (WCL) and fractionated conditioned media (CM) collected from serum starved HEK293Ts treated with vehicle or 20 nM BafA1 for 16 h. CM was subjected to serial differential ultracentrifugation to recover large EVs (10,000g; 10K), small EVPs (100,000g; 100K) and precipitated free soluble protein (TCA). Equal amounts of protein from WCL and fractionated CM were probed for the indicated proteins (n = 3). **(B)** Quantification of LC3 and autophagy cargo receptors in the indicated fractions of CM from serum starved BafA1 treated cells relative to WCL (mean ± s.e.m.; n = 3; p<0.05=*, p<0.01=**, p<0.005=***). Statistical significance between the CM fractions was calculated by one-way ANOVA with Tukey’s post hoc test. **(C)** Small EVPs from CM separated via linear sucrose density gradient ultracentrifugation, fractionated and immunoblotted to detect LC3, autophagy cargo receptors and the EV marker protein CD9 (n = 3). **(D)** Percent of total secreted LC3, cargo receptors and CD9 detected in individual linear sucrose gradient fractions (mean ± s.e.m.; n = 4). **(E)** Representative immunoblots of the indicated proteins from untreated EVPs or EVPs incubated with 100 µg ml−1 trypsin and/or 1% Triton X-100 (TX-100) for 30 min at 4 °C (n = 3). **(F)** Percent of protease protection for indicated proteins in EVPs incubated with 100 µg ml−1 trypsin and/or 1% Triton X-100 (mean ± s.e.m.; n = 3; ns=not significant, p<0.05= *, p<0.01= **, p<0.001=****).

### Autophagy cargo receptor secretion functionally requires multiple steps in classical autophagosome formation

The secretion of autophagy cargo receptors during lysosomal inhibition may be mediated by vesicular intermediates derived from classical double-membrane autophagosomes or proceed through emerging autophagy-related pathways that are independent of autophagosomes. To distinguish between these possibilities, we evaluated LC3 and autophagy cargo receptor secretion using a panel of ATG deficient cells that individually disrupt discrete steps required for autophagosome formation. Importantly, these analyses revealed that components required for phagophore formation (FIP200/RB1CC1, ATG14), LC3-conjugation (ATG7, ATG12) and the sealing of the double-membrane autophagosome (ATG2A/B) were all critical for the secretion of autophagy cargo receptors in response to BafA1 (Fig. 5A, B and Sup. Fig. 4A). Thus, in contrast to the previously described LDELS pathway, autophagosome formation is strictly required for the delivery of LC3 and cargo receptors outside the cell during lysosome impairment. In further support of this concept, BafA1-treated cells deficient for ATG14, which are defective for both autophagosome formation and cargo receptor secretion, showed significantly reduced co-localization of LC3 and p62/SQSTM1 with the late endosome and EV marker CD63 compared to controls (Fig. 5C,D and Sup. Fig. 4B). These results demonstrate that all of the key steps required for double-membrane autophagosome formation are functionally required for the capture and secretion of autophagy cargo receptors during lysosome impairment; furthermore, this material may transit through late endosomes along its itinerary to the extracellular space.

**Figure 5.**
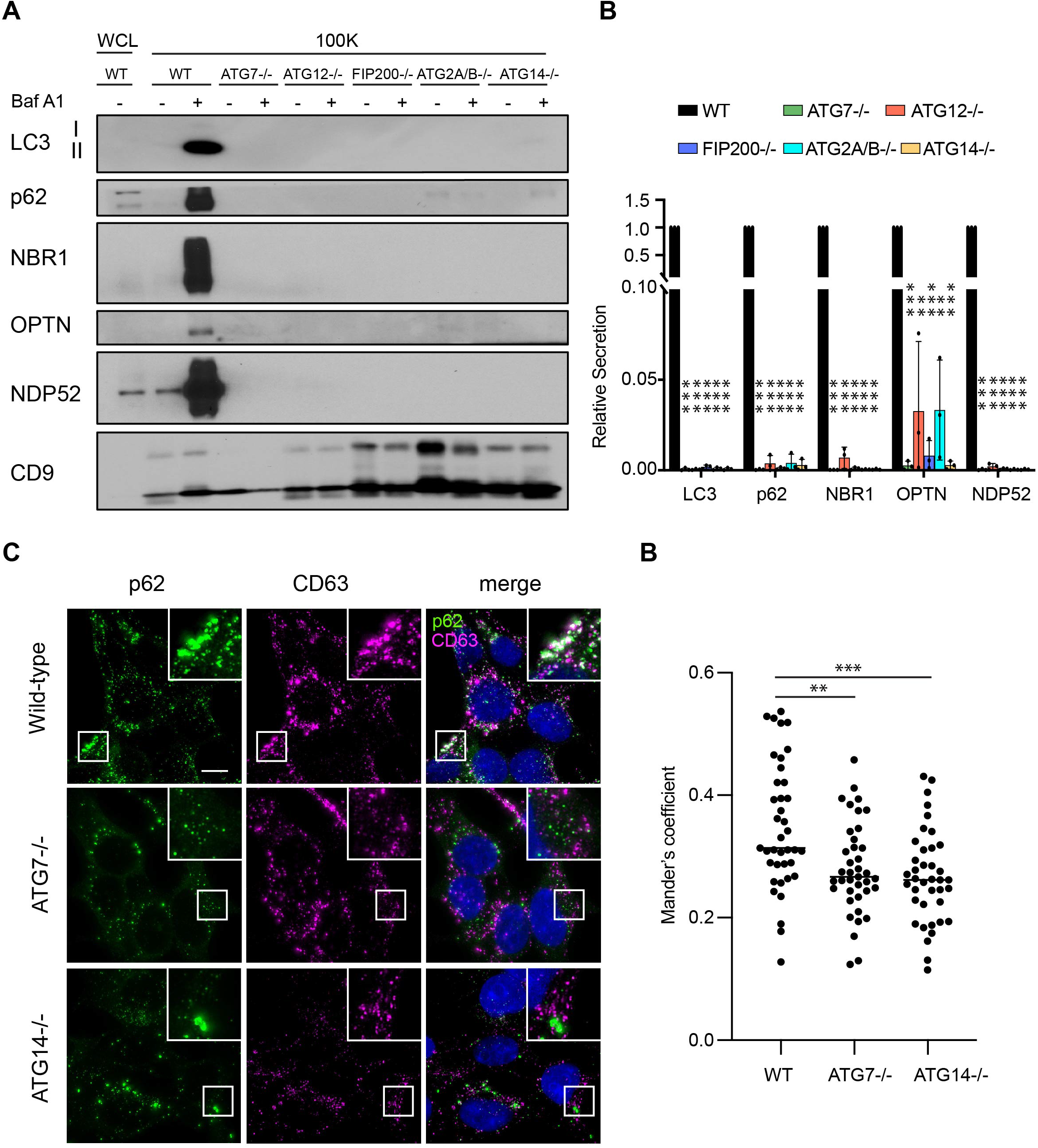
Autophagy cargo receptor secretion during lysosome inhibition requires autophagosome formation. **(A)** Whole cell lysate (WCL; left) and 100,000g EV fractions (100K; right) from serum starved wild-type (WT) and ATG-deficient HEK293Ts treated with vehicle or 20 nM nM BafA1 for 16h were collected, normalized to cell number and immunoblotted to detect LC3, autophagy cargo receptors and CD9 (n = 4). **(B)** Quantification of the levels of the indicated proteins in EVP fractions from BafA1 treated cells relative to treated WT controls. Statistical significance was calculated by one-way ANOVA coupled with Tukey’s post hoc test (mean ± s.e.m.; n=4; p<0.005=***). **(C)** Representative fluorescence micrographs from serum starved wild-type, ATG7−/− and ATG14−/− cells treated with 20 nM BafA1 and immunostained for endogenous p62 (green) and CD63 (magenta). Scale bar, 10 µm. **(D)** A scatter plot of Mander’s coefficients for the co-occurrence of p62 and CD63 in the immunostained cells in C. Statistical significance was calculated by an unpaired two-tailed t-test (mean ± s.e.m.; WT n = 39; ATG7−/− n=39, ATG14−/−, n = 39; p<0.01=**, p<0.01=***).

Evidence supports that lysosomal inhibitors perturb autophagosomes maturation into autolysosomes, resulting in the impaired turnover of autophagic material, which may subsequently be diverted for secretion outside the cells (Mauthe et al., 2018). Since our results with BafA1treatment demonstrated that autophagosomes are required for autophagy cargo receptor release, we interrogated whether the genetic disruption of autophagosome maturation was able to specifically promote this secretory phenotype. We evaluated autophagy cargo receptor secretion in cells deficient for SNAP29 or VAMP8, two SNAREs required for autophagosome-lysosome fusion (Itakura et al., 2012) (Fig 6A, B). Consistent with prior studies, genetic depletion of SNAP29 or VAMP8 led to impaired autophagosome maturation, evidenced by the striking accumulation of yellow autophagic puncta (mCherry-positive, GFP-positive) within cells that co-expressed the mCherry-GFP-LC3 flux reporter and intracellular accumulation of autophagy cargo receptors (Fig. 6C, D; Suppl. Fig. 5A,B). In addition to these impairments in autophagosome maturation, we observed increased LC3-II and autophagy cargo receptors secreted in the EVP fractions isolated from the conditioned media of SNAP29 and VAMP8 deficient cells compared to controls (Fig. 6E-H). Thus, similar to pharmacological lysosomal inhibition, the genetic inhibition of autophagosome-lysosome fusion is sufficient to drive the secretion of autophagy cargo receptors via EVPs.

**Figure 6.**
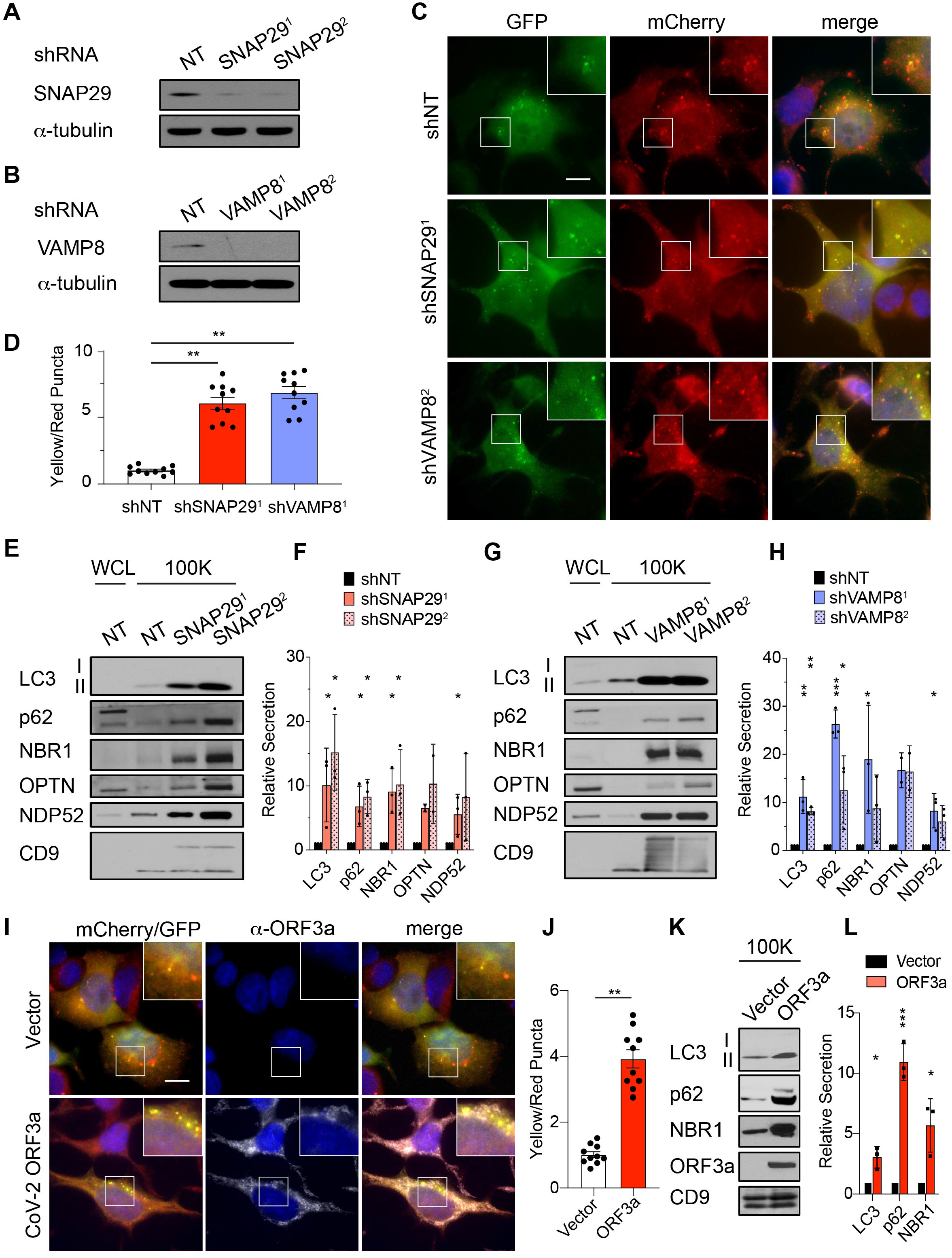
Genetic inhibition of autophagosome-lysosome fusion promotes the secretion autophagy cargo receptors with EVPs. **(A)** Lysate from cells stably expressing control shRNA (shNT) or shRNAs targeting SNAP29 (shSNAP29^1^; shSNAP29^2^) was blotted for SNAP29 and α-tubulin. **(B)** Lysate from cells that stably express control shRNA (shNT) or shRNAs targeting VAMP8 (shVAMP8^1^; shVAMP8^2^) was blotted for VAMP8 and α-tubulin. **(C)** Representative images of cells stably expressing the mCherry-EGFP-LC3 reporter and control shRNA (shNT), SNAP29 shRNA (shSNAP29^1^) or VAMP8 shRNA (shVAMP8^1^). Scale bar, 10 µm. **(D)** Quantification of the ratio of double-positive (mCherry+;EGFP+) yellow puncta to mCherry only (mCherry+;EGFP-) red puncta in cells stably co-expressing the mCherry-EGFP-LC3 reporter and the indicated shRNAs from C. (mean ± s.e.m.; shNT, n=10; shSNAP29^1^, n=10, shSNAP29^1^, n=10; p<0.01=**). Statistical significance was calculated by one-way ANOVA with Tukey’s post hoc test. **(E)** Whole cell lysate (WCL; left) and 100,000g EVP fractions (100K; right) from serum starved cells expressing control shRNA (NT) or SNAP29 shRNA (shSNAP29^1^; shSNAP29^2^) was blotted for the indicated proteins. **(F)** Quantification of the indicated proteins in EVPs from cells expressing SNAP29 shRNA relative to control shRNA in E. Statistical significance was calculated by one-way ANOVA with Tukey’s post hoc test (mean ± s.e.m.; n=3; p<0.05=*). **(G)** Whole cell lysate (WCL; left) and 100,000g EV fractions (100K; right) from serum starved cells expressing control shRNA (NT) or VAMP8 shRNA (shVAMP8^1^; shVAMP8^2^) was blotted for the indicated proteins. **(H)** Quantification of the indicated proteins in EVPs from cells expressing VAMP8 shRNA relative to control shRNA in G. Statistical significance was calculated by one-way ANOVA with Tukey’s post hoc test (mean ± s.e.m.; n = 3; p<0.05=*, p<0.01=**, p<0.005=***). **(I)** Representative images of cells that stably co-express the mCherry-EGFP-LC3 reporter SARS-CoV-2 ORF3a (2xStrepTag) or vector controls stained with anti-StrepTag antibody. Scale bar, 10 µm. **(J)** Quantification of the ratio of double-positive (mCherry+;EGFP+) yellow puncta to mCherry only (mCherry+;EGFP-) red puncta in cells stably co-expressing the mCherry-EGFP-LC3 reporter and SARS-CoV-2 ORF3a or vector controls in I. Statistical significance was calculated by unpaired two-tailed t-test (mean ± s.e.m.; vector, n = 10; SARS-CoV-2 ORF3a, n = 10; p<0.01=**). **(K)** 100K fractions from serum starved cells expressing SARS-CoV-2 ORF3a (2xStrepTag) or vector control were collected and blotted for the indicated proteins. **(L)** Quantification of the levels of LC3, p62 and NBR1 in EVPs from cells stably expressing SARS-CoV-2 ORF3a or vector in K. Statistical significance was calculated by one-way ANOVA coupled with Tukey’s post hoc test (mean ± s.e.m.; n=3; p<0.05=*, p<0.005=***).

### SARS-CoV-2 ORF3a inhibits autophagosome-lysosome fusion and promotes cargo receptor secretion via EVPs

Multiple small RNA viruses, including SARS-CoV-2, impair autophagosome-lysosome fusion in order to evade clearance as well as to generate intracellular membranes important for viral replication and packaging (Miao et al., 2021; Zhang et al., 2021). Therefore, we tested a recently identified viral inhibitor of autophagosome maturation, SARS-CoV-2 ORF3a, for its impact on the secretion of autophagy cargo receptors. Consistent with recent reports, cells stably expressing ORF3a had marked accumulation of early autophagosomes relative to controls in mCherry-GFP-LC3 reporter assays, characteristic of impaired autophagosome maturation (Fig. 6I, J). Moreover, EVPs fractioned from the conditioned media secreted by ORF3a expressing cells were enriched in LC3-II, p62/SQSTM and NBR1; notably, these fractions also contained SARS-CoV-2 ORF3a itself (Fig. 6K, L). These results further corroborate that impaired autophagosome-to-autolysosome maturation promotes the diversion of autophagic vesicular intermediates containing LC3 and autophagy cargo receptors for EVP-mediated secretion.

### Rab27a promotes autophagy cargo receptor secretion via EVPs

A subpopulation of EVPs is formed at late endosomes termed multivesicular endosomes (MVEs). Autophagosomes are also known to deliver material to MVEs, suggesting that LC3 and autophagy cargo receptors may be diverted through MVEs when autophagosome maturation into autolysosomes is impaired to facilitate extracellular release. Rab27a, a small GTPase that promotes the fusion of endolysosomal intermediates with the plasma membrane, has been implicated in multiple regulated lysosomal and granule exocytosis pathways, including the extracellular release of EVPs originating from MVEs (Desnos et al., 2003; Ostrowski et al., 2010; Stinchcombe et al., 2001). Thus, we tested whether Rab27a was functionally required for autophagy cargo receptor secretion; indeed, stable Rab27a knockdown suppressed the secretion of LC3, p62, NBR1, OPTN and NDP52 via EVPs in BafA1-treated cells (Fig. 7 A, B). Importantly, assays of autophagic flux confirmed that lyososomal turnover of LC3-II and p62/SQTM1 was unaltered in Rab27a deficient cells, indicating that Rab27a is specifically required for the secretion of cargo receptors in response to BafA1, rather than modulating the classical autophagic degradation of these components (Fig. 7C). Altogether, these results demonstrate that Rab27a is functionally required for the secretion of LC3 and cargo receptors in EVP populations. Moreover, these results broach that during lysosome impairment, early autophagic intermediates are diverted to MVEs requiring Rab27a-dependent exocytosis for extracellular release.

**Figure 7.**
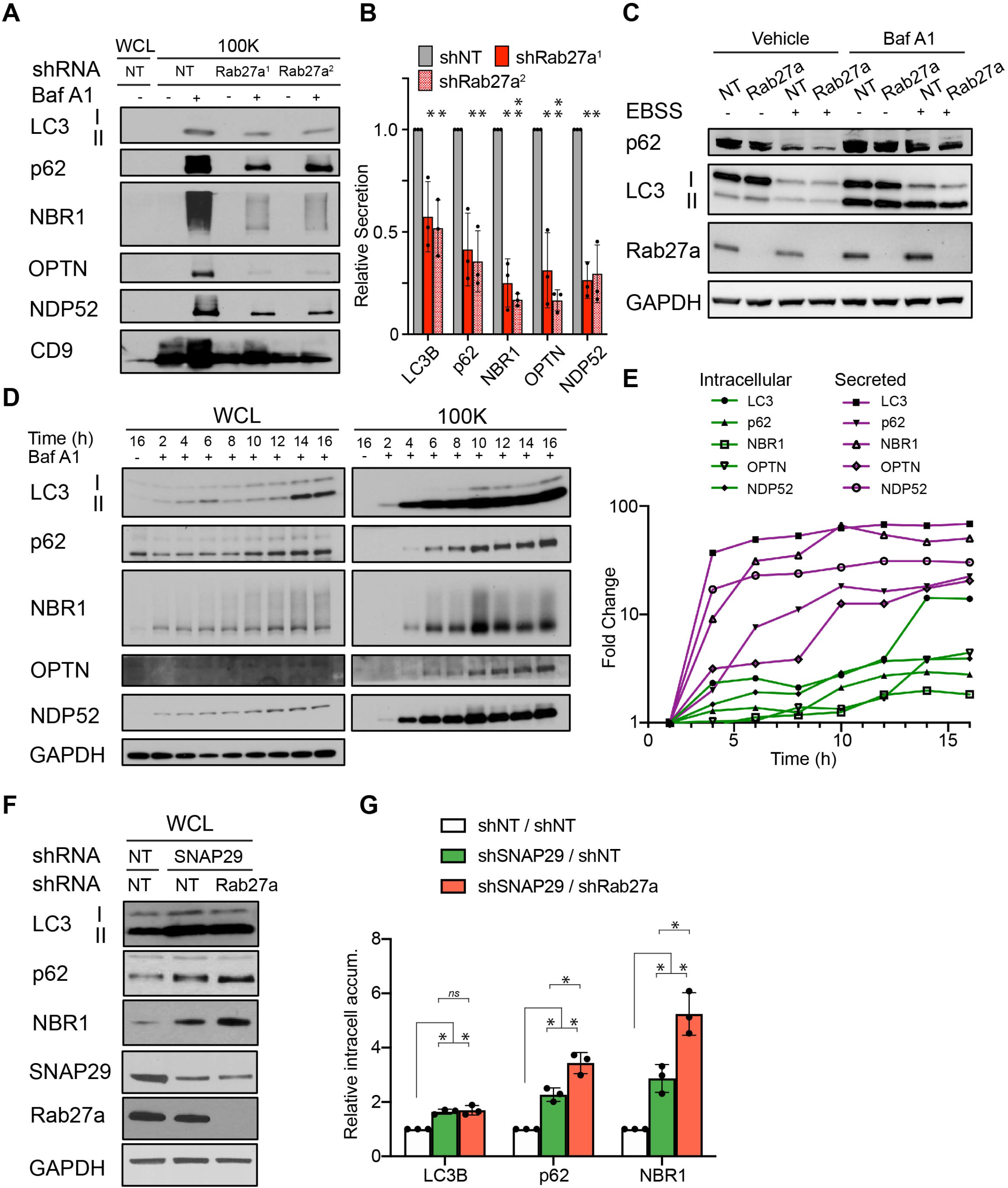
Rab27a is required for EVP secretion of autophagy cargo receptors in response to lysosome inhibition and impaired autophagosome maturation. **(A)** Whole cell lysate (WCL; left) and 100,000g EVP fractions (100K; right) from serum starved cells expressing control shRNA (NT) or Rab27a shRNA (shRab27a^1^; shRab27a^2^) were collected and blotted for the indicated proteins. **(B)** Quantification of the indicated proteins in EVPs from cells expressing SNAP29 shRNA relative to control shRNA in A. Statistical significance was calculated by one-way ANOVA coupled with Tukey’s post hoc test (mean ± s.e.m.; n=3; p<0.05=*, p<0.05=**). **(C)** Cells that express control shRNA (NT) or Rab27a shRNA (shRab27a1) were EBSS starved for 12 h, lysed and immunoblotted for the indicated proteins. Baf A1=50 nM BafA1 for 1 h before lysis (n=3). **(D)** WCL (left) and 100K fractions (right) from serum starved cells treated with 20 nM BafA1 were collected at the indicated times post-treatment and immunoblotted for LC3, autophagy cargo receptors and GAPDH. **(E)** Quantification of the fold change in intracellular levels (intracellular; shapes with green lines) and EVP-mediated secretion (secreted; shapes with magenta lines) for the indicated proteins relative to levels at 2 h post-treatment with BafA1 in D. **(F)** WCL from cells stably expressing control shRNAs (NT), SNAP29 shRNA and control shRNA (NT), or SNAP29 shRNA and Rab27a shRNA was collected 8 h post-starvation and blotted for the indicated proteins. **(G)** Quantification of LC3, p62 and NBR1 in WCL from starved cells expressing SNAP29 shRNA and control shRNA (shSNAP29/shNT) or SNAP29 and Rab27a shRNA (shSNAP29/shNT) relative to shRNA controls (shNT/shNT) in F. Statistical significance was calculated by one-way ANOVA coupled with Tukey’s post hoc test (mean ± s.e.m.; n = 3; ns=not significant, p<0.05=*).

### Secretory autophagy reduces intracellular accumulation of autophagy cargo receptors when degradative autophagy is impaired

In the setting of autophagy or lysosomal deficiency, the aberrant intracellular accumulation of autophagy cargo receptors, including p62/SQSTM1 and NBR1, is associated with numerous detrimental effects ranging from perturbations in intracellular signaling, changes in cellular differentiation, and the formation of intracellular inclusions (Bjorkoy et al., 2005; Mancias and Kimmelman, 2016; Marsh et al., 2020). Because cells may control the intracellular levels of these proteins through multiple routes, including extracellular secretion, we postulated that the diversion of autophagy cargo receptors for extracellular secretion via EVPs serves as a mechanism to control intracellular accumulation of these important effector proteins when autophagosome maturation and lysosomal degradation are impaired. To test this hypothesis, we first assayed the kinetics of autophagy cargo receptor secretion and accumulation in cells following acute treatment with BafA1 over 16h. Notably, the secretion of autophagy cargo receptors in EVP fractions was rapidly induced within 4h of BafA1 treatment and continued to increase until approximately 8-10h, at which time it reached a steady state (Fig. 7D, E, magenta lines). In contrast, intracellular accumulation of LC3 and cargo receptors was initially modest from 2-10h post-treatment; thereafter, more robust intracellular accumulation of these targets was observed at approximately 10-16h of lysosome inhibition (Fig. 7D, E, green lines). These observations highlighted that BafA1-induced extracellular secretion of autophagy cargo receptors precedes their robust intracellular accumulation, and suggested that secretory autophagy via EVPs buffers against the intracellular accumulation of autophagy cargo receptors when autophagy-dependent degradation is impaired.

To determine whether secretory autophagy directly impacts the intracellular accumulation of autophagy cargo receptors in response to impaired autophagy-dependent degradation, we scrutinized the effects of the combined genetic inhibition of degradative and secretory autophagy on intracellular levels of p62 and NBR1. Indeed, our results above demonstrated that genetic depletion of SNAP29 impaired autophagosome maturation, resulting in increased autophagy cargo receptor secretion. In contrast, Rab27a was functionally required for cargo receptor secretion in response to lysosomal inhibition, but had minimal effect on degradative autophagy. Accordingly, if secretory autophagy is important for mitigating the effects of autophagy-dependent cargo receptor degradation, cells doubly deficient for SNAP29 and Rab27a would be predicted to intracellularly accumulate autophagy cargo receptors at increased levels. To test this prediction, we compared intracellular cargo receptor levels in cells genetically deficient for SNAP29 versus those doubly deficient for SNAP29 and Rab27a. Cells genetically depleted of SNAP29 and Rab27a demonstrated significantly increased intracellular accumulation of p62/SQSTM1 and NBR1 relative to cells depleted of SNAP29 alone or wild type controls (Fig. 7 F,G). Thus, the secretion of autophagy cargo receptors via EVPs serves as an important homeostatic mechanism for balancing the intracellular levels of these autophagy targets in response to impaired autophagosome maturation and lysosomal degradation.

## Discussion

Here, we delineate a secretory autophagy process induced during lysosomal inhibition that targets autophagic cargo receptors for release via EVP-associated secretion pathways when autophagy-dependent degradation is impaired. Previous work has revealed important interconnections between autophagy and EV secretion. For example, our group has demonstrated that LC3-conjugation components regulate the loading and release of small EVs (Leidal et al., 2020). In addition, infectious pathogens and pharmacological agents that disrupt endolysosome function and autophagic turnover also facilitate the secretion of autophagy components and autophagy cargo receptors in EVs (Hessvik et al., 2016; Miao et al., 2015; Miranda et al., 2018). These connections have led both us and others to conjecture that autophagy and EVP secretion pathways function cooperatively to maintain cellular homeostasis (Leidal and Debnath, 2021; Xu et al., 2018). In this study, we illuminate that impaired autophagosome maturation promotes the rerouting of autophagic material through late endosomal intermediates, which results in its secretion in EVP-associated fractions via Rab27a-dependent exocytosis.

Importantly, this pathway of secretory autophagy during lysosomal inhibition is distinct from LDELS, which specifies the loading of RNA-binding proteins into intraluminal vesicles and their subsequent secretion within EVs (Leidal et al., 2020). The LDELS pathway is one of a growing number of recently identified autophagy-related processes, including LC3-associated phagocytosis (LAP) and LC3-assocated endocytosis (LANDO), in which the autophagy conjugation machinery mediates the conjugation of LC3 to single membranes organelles of the endolysosomal system via pathways distinct from classical autophagy (Nieto-Torres et al., 2021). In contrast, autophagy cargo receptor secretion in response to impaired lysosome function or autophagosome maturation is more closely akin to classical autophagy in that it requires multiple ATGs that mediate the progressive steps in autophagosome formation. Overall, our results motivate the model that defects in autophagosome-to-autolysosome maturation promote the accumulation of immature autophagosomes, which fuse with late endosomes and multivesicular bodies (MVBs), forming amphisomes. As a result, autophagic material such as autophagy cargo receptors can mix with intraluminal vesicles and particles and this material is subsequently released with EVPs upon amphisome fusion with the plasma membrane via a Rab27a-dependent process. In support, we observe significantly reduced intracellular co-localization of LC3 and p62/SQSTM1 with the late endosome marker CD63 in ATG14 deficient cells in comparison to wild type controls, when both are treated with BafA1.

We propose that secretory autophagy during lysosomal inhibition utilizes EVP-based secretion pathways as an alternate route to clear sequestered material when degradation is impaired. As a result, blocking Rab27a-mediated exocytosis not only reduces the extracellular secretion of autophagy cargo receptors but also exacerbates the accumulation of these proteins when autophagosome-to-autolysosome maturation is simultaneously impaired. Defects in autophagic flux and lysosomal function are observed both during aging and various age-related disorders and secretory autophagy may serve particularly important roles in maintaining proteostasis in these settings, especially in non-dividing cells such as neurons (Leidal et al., 2018). An exciting question for future study is determining whether lysosomal dysfunction in age-related neurodegenerative disorders, such as Parkinson disease (PD), Alzheimer disease (AD) and Frontotemporal Dementia (FTD), elicits the increased secretion of autophagy cargo receptors, and if so, scrutinizing how secretory autophagy impacts proteostasis in such patients (Leidal et al., 2018). Nevertheless, despite these potential cell-autonomous benefits of secretory autophagy in mitigating intracellular protein accumulation during lysosomal inhibition, the extracellular secretion of autophagy cargo receptors may harbor deleterious effects *in vivo*. Notably, recent work demonstrates that extracellular p62/SQSTM1 promotes sepsis-induced death in mice due to disruptions in immune metabolism and increased systemic coagulation (Zhou et al., 2020). Hence, further understanding whether and how the EVP secretion of p62/SQSTM1 and other cargo receptors modulates inflammatory and immune function in various human pathologies due to defective lysosome function remains an important question for future study. Interestingly, we demonstrate that the SARS-CoV2 ORF3a viral product not only disrupts autophagosome-to-autophagosome maturation but potently promotes the extracellular secretion of p62/SQSTM1 via EVPs (Miao et al., 2021; Zhang et al., 2021). Accordingly, further scrutinizing whether and how extracellular p62/SQSTM1 secreted from SARS-CoV2-infected cells contributes to COVID19-associated immune pathologies and coagulopathies remains an important future direction (Fajgenbaum and June, 2020).

Finally, we show that treatment with the antimalarial chloroquine (CQ) in vivo results in the increased EVP secretion of autophagy cargo receptors in murine plasma. Over the past several years, hydroxychloroquine (HCQ) has been actively pursued to therapeutically inhibit the autophagy-lysosome pathway in human oncology trials (Amaravadi et al., 2019). However, a critical barrier for assessing the efficacy of such strategies has been non-invasively monitoring changes in autophagic flux in humans, particularly in response to the therapeutic modulation of autophagy and lysosomal pathways (Mizushima and Murphy, 2020). Given the continued interest in targeting autophagy and lysosomal pathways for cancer treatment combined with the growing excitement regarding EVPs proteins as non-invasive liquid biopsy tools, measuring the autophagy-dependent EVP secretion of cargo receptors in human plasma potentially represents a powerful biomarker for monitoring the efficacy of next-generation lysosomal inhibitors in cancer treatment, and more broadly, for monitoring lysosomal dysfunction in diverse therapeutic settings.

## Acknowledgements

Grant support includes the NIH (CA201849, CA126792, CA213775 to JD, AG057462 to JD and EJH, CA226851 to APW, and F31CA217015 to TM), Samuel Waxman Cancer Research Foundation (to J.D.), Mark Foundation for Cancer Research (Endeavor Award to JD), UCSF QB3 Calico Longevity Fellowship (to JD and AML) and Dale Frey Breakthrough Award from the Damon Runyon Cancer Research Foundation (DFS 14-15 to APW). Fellowship support includes a Banting Postdoctoral Fellowship from the Government of Canada (201409BPF-335868) and Cancer Research Society Scholarship for Next Generation of Scientists (22805) to AML, and a NSF Graduate Student Fellowship (1650113 to TAS).

## Author contributions

JD, AML, EJH and TAS conceived the study and designed the experiments. TAS, TAN, YTL, APW, and JD designed and optimized the quantitative proteomics workflows. TAS, TAN, YTL and performed mass spectrometry and analyzed the resulting LC-MS/MS data with input from AML, APW, EJH and JD. TAS, TAN, AML, performed biochemical and cell biological experiments. TAS, AML and TM performed mouse experiments. AML, TAS and JD wrote the paper, with input from all other authors. JD, EJH and AML supervised the study.

## Methods

### Cell culture

HEK-293T (ATCC, CRL-3216), MDA-MB-231 (ATCC, CRM-HTB-26) and R221A cells (Gift from Barbara Fingleton) were cultured in Dulbecco’s Modified Eagle Medium (DMEM), high glucose, pyruvate (Gibco 11995040) supplemented with 10% fetal bovine serum (FBS) (Atlas Biologicals F-0500-D), 25 mM HEPES (Teknova H1030), 100 units/mL penicillin and 100 μL/mL streptomycin (Gibco 14140163). All cell lines were tested for mycoplasma contamination (Sigma-Aldrich MP0035).

### CRISPR/Cas9 gene-deletion

HEK-293T knockout cell lines were generated by transient transfection of pSpCas9(BB)-2A-Puro (Addgene 48139) encoding U6 driven expression of sgRNAs (Scramble Guide: 5’-GCACTACCAGAGCTAACTCA-3’; ATG7 Guide: 5’-ACACACTCGAGTCTTTCAAG-3’; ATG12 Guide: 5’-CCGTCTTCCGCTGCAGTTTC-3’; ATG14 Guide: 5’-CTACTTCGACGGCCGCGACC-3’; FIP200 Guide: 5’-AGAGTGTGTACCTACAGTGC-3’; ATG2A Guide: 5′-CGCTGCCCTTGTACAGATCG-3′; ATG2B Guide: 5′-ATGGACTCCGAAAACGGCCA-3′). Cells were selected 48-72 hours post-transfection with 1 g/ml puromycin for 48 h. Polyclonal populations were collected for Surveyor analysis (IDT, 706020) and were sorted into single-cell populations by limiting dilution at 1.5 cells/well per 96-well plate. For DNA analysis, genomic DNA samples were prepared using QuickExtract (Epicentre). The PCR products were column purified and analyzed with Surveyor Mutation Detection Kit (IDT). For genotyping of single-sorted cells, PCR amplified products encompassing the edited region (ATG7 Fwd: 5’-TGGGGGACAGTAGAACAGCA-3’, ATG7 Rev: 5’-CCTGGATGTCCTCTCCCTGA-3’; ATG12 Fwd: 5’-AGCCGGGAACACCAAGTTT-3’, ATG12 Rev: 5’-GTGGCAGCCAAGTATCAGGC-3’; ATG14 Fwd: 5’-AAAATCCCACGTGACTGGCT-3’, ATG14 Rev: 5’-AATGGCAGCAACGGGAAAAC-3’; FIP200 Fwd: 5’-ATTCTCTGGCTTGACAGGACAG-3’, FIP200 Rev: 5’-AAATACTGAGCGTGCACATTGC-3’; ATG2A Fwd: 5’-GAGCCGGACGGGGATCGC-3’, ATG2A Rev: 5’-CTGCAAGGTGAGCTGGAGGC-3’; ATG2B Fwd: 5’-ATAGGGATGGAGGGGCCGC-3’, Rev: 5’-TCATTGAGACACCATTTGTC-3’ were cloned into pCR™4-TOPO® TA vector using the TOPO-TA cloning kit (Thermo Fisher 450030) and sequence verified.

### Plasmid Constructs

The following vectors are available or were obtained from Addgene: pBABE-puro mCherry-EGFP-LC3 (#22418), pBABE-hygro (#1765), psPAX2 (#12260), pMD2.G (Plasmid #12259), pLKO.1-blast (#26655) and pLVX-EF1alpha-SARS-CoV-2-orf3a-2xStrep-IRES-Puro(#141383). pBABE-hygro mCherry-EGFP-LC3 was generated by subcloning mCherry-EGFP-LC3 from pBABE-puro mCherry-EGFP-LC3 into shared AgeI and SalI restriction sites in pBABE-hygro. pLKO.1 lentiviral vectors for shRNAs targeting SNAP29 (#1 TRCN0000083650; #2 TRCN0000231850), VAMP8 (#1 TRCN0000219911; #2 TRCN0000181123), Rab27a (#1 TRCN0000005296; #2 TRCN0000005297), and non-targeting control (SHC002) are available from Sigma-Aldrich. pLKO.1-blast shSNAP29 was generated by subcloning the the blasticidin resistance gene from pLKO.1-blast into shared BamHI and KpnI restriction sites in pLKO.1 shSNAP29 (#TRCN0000083650).

### Retroviral and lentiviral packaging, infection and selection

Retroviral pBABE expression vectors were packaged and target cells were transduced according to established protocols. Briefly, Phoenix-AMPHO cells (gift from C. McCormick, Dalhousie University) were seeded and transfected with retroviral vectors using polyethylenimine. Virus-containing CM was collected 2 d after transfection and clarified using a 0.45-µM filter. Prior to infection, virus-containing medium was diluted 1:4 in DMEM growth medium and the mix was supplemented with Polybrene to a final concentration of 8 µg ml−1. Subsequently, the viral transduction mix (5 ml total volume/10 cm culture dish) was incubated with HEK293T cells for 24 h. Cells were selected 24 h post-transduction with 1 µg ml^−1^ puromycin for 2 d or 100 µg ml^−1^ hygromycin for 5 d. To package lentivirus, HEK293T cells were seeded and co-transfected with the packaging vectors psPAX2 and pMD2.G, and individual pLKO.1 transfer vectors or pLVX-EF1alpha-SARS-CoV-2-orf3a-2xStrep-IRES-Puro. Virus collection and infection was carried out as above. Stably transduced cell pools were selected with with 1 µg ml^−1^ puromycin for 2 d or 10 µg ml^−1^ hygromycin for 4 d.

### Antibodies

The following antibodies were used for immunoblotting: rabbit anti-ATG14 (MBL, PD026; 1:1,000), rabbit anti-LAMP1 (Cell Signaling, 9091; 1:1,000), rabbit anti-Rab7a (Cell Signaling, 9367; 1:1,000), rabbit anti-Rab5a (Cell Signaling, 3547; 1:1,000), mouse anti-GM130 (BD Biosciences, 610823; 1:1,000), rabbit anti-CYC1 (Novus Biologicals, NBP1-86872; 1:1,000), mouse anti-NUP62 (BD Biosciences, 610497; 1:1,000), mouse anti-GAPDH (Millipore, MAB374; 1:10,000), rabbit anti-LC3 (Millipore, ABC232; 1:1,000), guinea pig anti-p62/SQSTM1 (Progen, GP62-C; 1:1,000), rabbit anti-p62/SQSTM1 (Cell Signaling, 5114; 1:1,000), mouse anti-NBR1 (Santa Cruz Biotechnology, sc-130380; 1:500), rabbit anti-OPTN (Abcam, ab23666; 1:1,000), mouse anti-NDP52/CALCOCO2 (Abnova, H00010241-B01P; 1:1,000), mouse anti-TSG101 (BD Biosciences, 612696; 1:1,000), mouse anti-ALIX (Cell Signaling, 2171; 1:1,000), mouse anti-CD9 (Sigma-Aldrich, CBL162; 1:1,000), mouse anti-GFP (Santa Cruz sc-9996), rabbit anti-Rab27a (Cell Signaling, 69295; 1:1,000), rabbit anti-SNAP29 (Abcam, ab138500; 1:1,000), rabbit anti-VAMP8 (Abcam, ab76021; 1:1,000), rabbit anti-StrepTag II (Abcam, ab76949; 1:500) Peroxidase-AffiniPure Donkey Anti-Rabbit IgG (H+L) (Jackson, 711-035-152; 1:5,000), Peroxidase-AffiniPure Donkey Anti-Guinea Pig IgG (H+L) (Jackson, 706-035-148; 1:5,000), Peroxidase-AffiniPure Donkey Anti-Goat IgG (H+L) (Jackson, 705-035-147; 1:5,000), and Peroxidase-AffiniPure Donkey Anti-Mouse IgG (H+L) (Jackson, 715-035-150; 1:5,000).

For immunofluorescence we employed: rabbit anti-LC3B (1:200, Cell Signaling Technology, #3868T), mouse anti-CD63 (1:200, Abcam, #ab8219), guinea pig anti-p62/SQSTM1 (Progen/Cedarlane, #GP62-C; 1:1,000), rabbit anti-StrepTag II (Abcam, ab76949; 1:200), AlexaFluor goat anti-rabbit 488 (1:500, Thermo Fisher, #A-11034), goat anti-mouse 568 (1:500, Thermo Fisher, #A-11031) and goat-anti guinea pig (ThermoFisher A-21450, 1:1000).

### Immunoblotting

For whole cell lysate (WCL), cells were lysed in RIPA buffer (25 mM Tris-HCl pH 8.0, 150 mM NaCl, 1% NP-40, 1% sodium deoxycholate, 0.1% SDS) supplemented with 10 mM NaF, 10 mM β-glycerophosphate, 1 mM Na_3_VO_4_, 10 nM calcyulin A, 0.5 mM phenylmethyl sulphonyl fluoride, 0.1 mM E-64-D, 10 μg/mL pepstatin A, and protease inhibitor cocktail. Lysates were cleared by centrifugation at 15,000 g for 10 min at 4°C. For EV lysate, pelleted EVs were resuspended in urea buffer (50 mM Tris-HCl, pH 8.0, 8 M urea, 2% SDS, 10 mM NaF, 5 mM EDTA). WCL and EV lysates were quantified by BCA assay (Thermo Fisher 23225), mixed with sample buffer, resolved by SDS-PAGE, and transferred to polyvinylidene fluoride membrane. Membranes were then blocked in 5% milk in PBS with 0.1% Tween 20 (PBST), incubated in primary antibody overnight at 4°C in blocking buffer, washed with PBST, incubated in HRP-conjugated secondary antibodies for 1 h at room temperature in blocking buffer, washed with PBST, and visualized by enhanced chemiluminescence (Thermo Fisher 32106) on film. Immunoblots were quantified by densitometry using Fiji (ImageJ v.2.0.0-rc-69/1.52p).

### Fluorescence and immunofluorescence microscopy

To perform mCherry-GFP-LC3 fluorescence microscopy, cells were stably transduced with retroviral vectors that encoding mCherry-GFP-LC3 and puromycin or hygromycin resistance genes for 24h and selected in 1 μg/mL puromycin or 100 μg/mL hygromycin B (ThermoFisher #10687010) for two or five days, respectively. Subsequently, cells were seeded onto coverslips coated with 10 μg/mL fibronectin and 24h later treated with vehicle or 20 nM BafA1 in serum-free DMEM for 16 h. Cells were fixed with 4% PFA in PBS for 15 min at room temperature, washed with PBS, and mounted onto slides with Prolong Gold Antifade with DAPI (Thermo Fisher P36935). Cells were visualized using a DeltaVision microscope (Applied Precision Ltd.) fitted with a 60Å∼, 1.4-NA objective and CoolSnap HQ camera (Photometrics). Images were acquired using softWoRx software (Applied Precision Ltd.) and prepared in Fiji and Adobe Photoshop. To quantify mCherry-EGFP-LC3 puncta, cells were outlined manually, and each channel was independently autothresholded using the same settings across all images from each channel. mCherry and GFP puncta were then quantified using the Analyze Particles plugin in ImageJ. Double-positive puncta were identified and counted using the Colocalization and Analyze Particles plugins in ImageJ.

For immunofluorescence of endogenous p62 and CD63 or LC3 and CD63, cells were seeded on coverslips coated with 10 µg/mL fibronectin and 24h later treated with vehicle or 20 nM BafA1 in serum-free DMEM for 16 h. Subsequently, cells were fixed with 4% PFA in PBS for 15 min at room temperature, permeabilized with ice-cold methanol and incubated at −20 °C for 5 min before quenching with PBS/glycine. Samples were then blocked in blocking buffer (PBS + 0.1% Tween + 10% goat serum) for 1 h at room temperature, incubated with guinea pig anti-p62 (and mouse anti-CD63 (1:200, Abcam ab8219) or rabbit anti-LC3B (1:200, MBL PM036) and mouse anti-CD63 (1:200, Abcam ab8219) antibodies for primary staining. Cells were visualized using a DeltaVision microscope (Applied Precision) fitted with a 60X, 1.4-NA objective and a CoolSnap HQ camera (Photometrics). Images were acquired using softWoRx software (Applied Precision) and prepared in Fiji and Adobe Photoshop. Costes significance tests for co-occurrence and the Mander’s overlap coefficient for p62 and CD63 or LC3 and CD63 were performed by drawing a region of interest around individual cells and then employing the Coloc 2 analysis function within Fiji (PSF: 20, Costes randomizations: 10).

### Cell viability assay

To quantify cell viability, cells were seeded in 12-well plates, and after 24 h were treated with vehicle, 20 nM BafA1 or 25 μM CQ in serum-free DMEM for 8h. Cells were collected and stained with 0.4% trypan blue (Thermo Scientific T10282). Live and dead cells were then enumerated using the Countess II Automated Cell Counter (Thermo Scientific AMQAX1000).

### Extracellular vesicle preparation and characterization

EVs were isolated according to standard differential centrifugation protocol as described previously^63^. Briefly, HEK-239T cells were seeded in 15 cm tissue culture dishes and, when confluent, incubated with serum-free DMEM +/− treatment (20 nM BafA1, 25 μM CQ) for 16 or 24h. Conditioned media was then collected and serially centrifuged: 200 g for 10 min at 4°C to pellet cells, 2,000 g for 10 min at 4°C to pellet cellular debris and apoptotic bodies, 10,000 g for 30 min at 4°C in an ultracentrifuge to pellet large EVs, and 110,000 g for 70 min or 3h at 4°C in an ultracentrifuge to pellet small EVs. The 110,000 g pellet was then gently resuspended in 1 mL PBS by pipetting, diluted in 12 mL PBS, and ultracentrifuged at 110,000 g for 70 min at 4°C. The final EV pellets were resuspended in urea lysis buffer for immunoblotting analysis, 10% sucrose for sucrose density gradient analysis, or PBS for protease protection analysis. For analysis of secreted soluble protein, the supernatant from the first 110,000 g spin was collected and precipitated by addition of 15% trichloroacetic acid (TCA), then incubated for 1 h on ice before centrifugation at 180,000 g for 2h at 4°C. The 180,000 g pellet was then resuspended in 10 mL ice-cold 100% acetone, centrifuged at 180,000 g for 1 h at 4°C, and finally resuspended in urea lysis buffer. For comparison of EVs between experimental conditions *in vitro*, EV quantification or EV protein quantification were corrected as indicated in figure legends based on total cell number or WCL protein quantification to correct for cell seeding differences between conditions.

Sucrose density gradient separation was used to analyze co-fractionation between LC3, ATG cargo receptors, and canonical EV markers. 10% and 60% sucrose solutions in PBS were used to prepare continuous 10-60% sucrose gradients on a gradient station (Biocomp Instruments 153). EV pellets were generated by differential centrifugation as described previously, then resuspended in 100 μL 10% sucrose solution, gently layered on top of the 10-60% gradient, and ultracentrifuged at 210,000 g for 18 h at 4°C. After ultracentrifugation, 1 mL fractions of the gradient were carefully top-unloaded, weighed, and diluted in 12 mL PBS. The diluted sucrose gradient fractions were ultracentrifuged at 110,000 g for 70 min at 4°C and pellets were resuspended in urea lysis buffer for immunoblotting analysis.

Protease protection analysis was used to determine membrane protection of secreted cargo receptors. EV pellets were generated by differential centrifugation as described previously, resuspended in 60 μL PBS, and divided equally into 3 fractions for resuspension in PBS, 100 μg/mL trypsin in PBS, or 1% NP-40 and 100 μg/mL trypsin in PBS. After incubation and occasional mixing for 30 min at 4°C, protease reactions were stopped with the addition of 10 μL protease inhibitor cocktail and 2x sample buffer before immunoblotting analysis.

Nanoparticle tracking analysis (NTA) technology was used to determine EV concentration and size. Conditioned media clarified of cell debris and large EVs by 0.22 μm filter were normalized by cell number, then loaded on a nanoparticle analyzer (Malvern Panalytical NanoSight NS300) and analyzed using NTA software (NTA 3.3 Dev Build 3.3.104). Camera level was set at 15 for all recordings. Camera focus was adjusted to make the particles appear as individual dots with surrounding refractory rings. Three 30 s videos were recorded for each sample with a delay of 5 s between each recording.

### Mass spectrometry of extracellular vesicle proteins

WT and *ATG7*-/- HEK293T cells were treated with 20nM BafA1 or vehicle in serum-free media for 16 h. After 16 h, conditioned media was collected and EVs were isolated using the differential centrifugation protocol outlined previously. Pelleted EVs were lysed in 800 μL RIPA buffer (25 mM Tris-HCl pH 8.0, 150 mM NaCl, 1% NP-40, 1% sodium deoxycholate, 0.1% SDS) supplemented with 2% SDS, then sonicated with a probe sonicator at amplitude 8 for 10 pulses of 10 s each. Samples were then diluted in 4 mL ice-cold 100% acetone and incubated at −20°C overnight. Samples were spun in an ultracentrifuge at 200,000 g for 18 h at 4°C, the acetone was decanted, and the samples air dried for 1 h at 25°C before being stored at −80°C until further processing.

Three biological replicates of precipitated EVs from WT and *ATG7*-/- bafA1-treated cells were resuspended in 30 μL 6 M guanidinium-chloride, 100 mM Tris pH 8.0, 10 mM TCEP, 40 mM 2-chloroacetamide. EV proteins were denatured for 1 h at 37°C, then quantified with 660nm Protein Assay Reagent (Thermo Scientific 22660). Samples were then diluted six-fold in 150 μL 100 mM Tris pH 8.0. 125-150 μg of protein for each sample was trypsinized with 15 μg trypsin (Thermo Scientific 90057) in an orbital shaker at 250 rpm, 37°C for 20 h. Trypsin digestion was then stopped by adding 10% trifluoracetic acid (TFA) to a final concentration of 0.5% TFA. Samples were desalted with SOLA solid phase extraction cartridges (Thermo Scientific 60109). Briefly, columns were washed with 500 μL 100% acetonitrile (ACN), then equilibrated with 500 μL 0.1% TFA twice before adding sample. Samples were washed with 500 μL 0.1% TFA three times, then 500 μL 0.1% formic acid (FA), 2% ACN, and eluted with 450 μL 0.1% FA, 50% ACN. Samples were dried by speed-vac, then resuspended in 10 μL 50 mM HEPES pH 8.5 and quantified by Pierce Quantitative Colorimetric Peptide Assay (Thermo Scientific 23275).

For Tandem Mass Tag (TMT) labeling, 800 μg of TMT^10^-126, 127N, 128C, 129N, 130C, and 131 (Thermo Scientific 90110) was reconstituted with 41 μL 100% anhydrous ACN. 15 μg of peptides from each replicate were individually combined with 7.69 μL (150 μg) of the TMT isobaric tags. Samples were incubated at 25°C for 1 h before quenching the reaction with 8 μL 5% hydroxylamine for 15 min. After labeling, the six individually labeled samples were pooled, dried by speed-vac, and then resuspended in 300 μL 0.1% TFA. The pooled samples were then fractionated into 8 fractions using Pierce High pH Reversed-Phase Peptide Fractionation Kit (Thermo Scientific 84868), then dried by speed-vac. The fractions were then resuspended in 0.1% FA, 2% ACN before LC-MS/MS analysis.

For LC-MS/MS analysis, 1 μg TMT-labeled peptide per fraction was analyzed on a 15 cm C18 analytical column, in-line with a Q-Exactive Plus mass spectrometer. The peptides were separated on a multi-slope, 100 min gradient (6.4% - 27.2% ACN with 0.1% FA for 80 min at 0.2 μL/min, then 27.2% - 40% ACN with 0.09% FA for 15 min at 0.3 μL/min, then 40% - 56% ACN with 0.09% FA for 5 min at 0.3 μL/min, and then washed for 3 min). Data dependent acquisition with MS1 resolution of 70,000, top15 method, and HCD normalized collision energy of 32 was used, with MS2 resolution of 35,000 and an isolation window of 0.7 m/z. Dynamic exclusion was activated for 30 s after initial parent ion selection.

Eight injections of the different fractions of TMT-labeled EV peptides were analyzed together via MaxQuant (v1.6.0.16). Search parameters for peptide search tolerance was 4.5 ppm, centroid match tolerance was 8 ppm, and 2 missed tryptic cleavages were permitted. Constant modification of carbamidomethylation of cysteines and variable modifications of N-terminal acetylation, methionine oxidation, and Ser/Thr/Tyr phosphorylation were allowed. Peptide spectrum match FDR and protein FDR were set at 1%. “Match between runs” was enabled to increase peptide identification. Type was set to “Reporter ion MS2,” and the six labels used in sample preparation were selected. The resulting quantifications were then median normalized for each channel, and statistical analysis (two-sample *t*-test) was applied in R with a statistical significance threshold of *p* < 0.05.

### Bioinformatic analyses

For mass spectrometry, the top eight proteins with the highest connectivity to candidates statistically enriched in EVs from wild-type cells treated with bafA1 versus EVs from *ATG7*-/- cells treated with bafA1 (p-value < 0.05; log_2_ WT/*ATG7-/-* > 0.585; n=3) were identified using the protein-protein interaction (PPI) hub protein tool in the Enrichr gene set enrichment analysis web server^64,65^ and plotted according to their −log_10_ adjusted p-values. GO analysis (database released 2019-12-09) of the secretome was performed with PANTHER Overrepresentation Test (released 2019-07-11)^66^ and the top ten terms for biological process and cellular component were plotted according to their −log_10_ false discovery rate. The overlaps between the defined secretome and the LDELS secretome (Leidal et al., 2020), ATG7 and ATG12-dependent EV secretome (Leidal et al., 2020), ATG5-dependent macrophage secretome (Kimura et al., 2017), autophagy interaction network (Behrends et al., 2010) and the autophagosome proteome from chloroquine treated cells (Mancias et al., 2014) were performed using Biovenn (Hulsen et al., 2008).

### In vivo extracellular vesicle isolation from mice

All experimental procedures and treatments were conducted in compliance with UCSF Institutional Animal Care and Use Committee (IACUC) guidelines under an approved animal protocol (#AN170608). C57BL/6JJc1 homozygous transgenic mice expressing GFP-LC3B^68,69^ (Riken Bioresource Center No. 00806) received either 60 mg/kg CQ or vehicle via intraperitoneal injection for 3 consecutive days. 6 h after final CQ injection, whole blood was collected in heparin-coated capillary tubes (Sarstedt CB 300 Lithium Heparin) and centrifuged at 2,000 g for 5 min at room temperature. The plasma phase was collected and 1 mL was pooled (from 3-4 mice per condition), diluted in 10 mL PBS, clarified by 0.22 μm filter, and ultracentrifuged at 110,000 g for 3 h at 4°C. The pellet was resuspended in 10 mL PBS, ultracentrifuged at 110,000 g for 70 min at 4°C, and resuspended in urea buffer for immunoblotting analysis. For comparison of EVs between experimental groups *in vivo*, samples were normalized by collection from equivalent volumes of plasma from experimental groups.

### Statistical analyses

Statistical analyses were performed using Prism GraphPad 5 software. Groups were compared using unpaired or paired Student’s t-test where indicated for pairwise comparisons or one-way ANOVA followed by Tukey’s post-hoc test for multiple comparisons. The sample size was chosen on the basis of the size of the effect and variance for the different experimental approaches. Details regarding the statistical analysis of the proteomic data and the bioinformatics analysis of the BafA1 induced ATG7-dependent EVP proteome are provided in the corresponding figure legends and/or the Methods sections above. P values of less than 0.05 are considered to be significant.

---

**Supplemental Figure 1.**
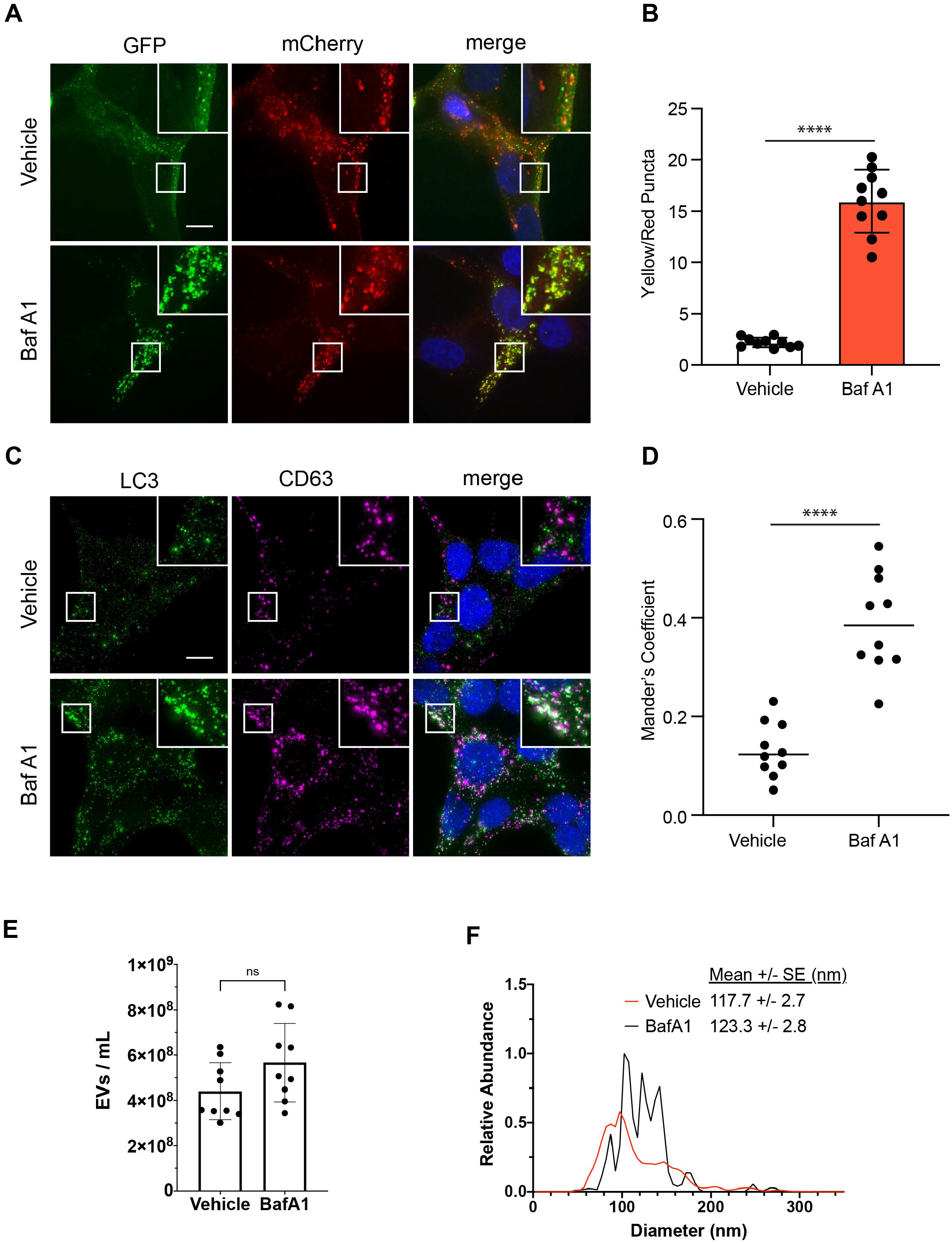
BafA1 treatment inhibits autophagic flux and modulates EV secretion. **(A)** Representative images of wild-type HEK293T cells stably expressing the mCherry-EGFP-LC3 reporter were treated with 20 nM BafA1 or vehicle (Vehicle) in serum-free media (BafA1) for 16 h. Scale bar, 10 µm. **(B)** Quantification of the ratio of double-positive (mCherry+/GFP+) to mCherry-only (mCherry+/GFP-) LC3 puncta per cell (mean +/− s.e.m.; vehicle, n=10; BafA1, n=10; p<0.005=***). **(C)** Representative images of wild-type HEK293T treated with 20 nM BafA1 or vehicle (Vehicle) in serum-free media (BafA1) for 16 h and immunostained for endogenous LC3 and CD63. Scale bar, 10 µm. **(D)** A scatter plot of Mander’s coefficients for the co-occurrence of LC3 with CD63 in the immunostained cells in C. The statistical significance was calculated by an unpaired two-tailed t-test (mean ± s.e.m.; Vehicle, n = 10; BafA1, n=10; p<0.005=***). **(E)** Nanoparticle tracking analysis of conditioned media from equal numbers of WT cells treated with vehicle in serum-free media (Vehicle) or 20 nM BafA1 (BafA1) (mean ± s.e.m.; n=3). Statistical significance calculated by unpaired two-tailed t-test (n.s.=not significant). **(F)** EV size distribution from indicated cell treatments in E (mean ± s.e.m.; n=3).

**Supplemental Figure 2.**
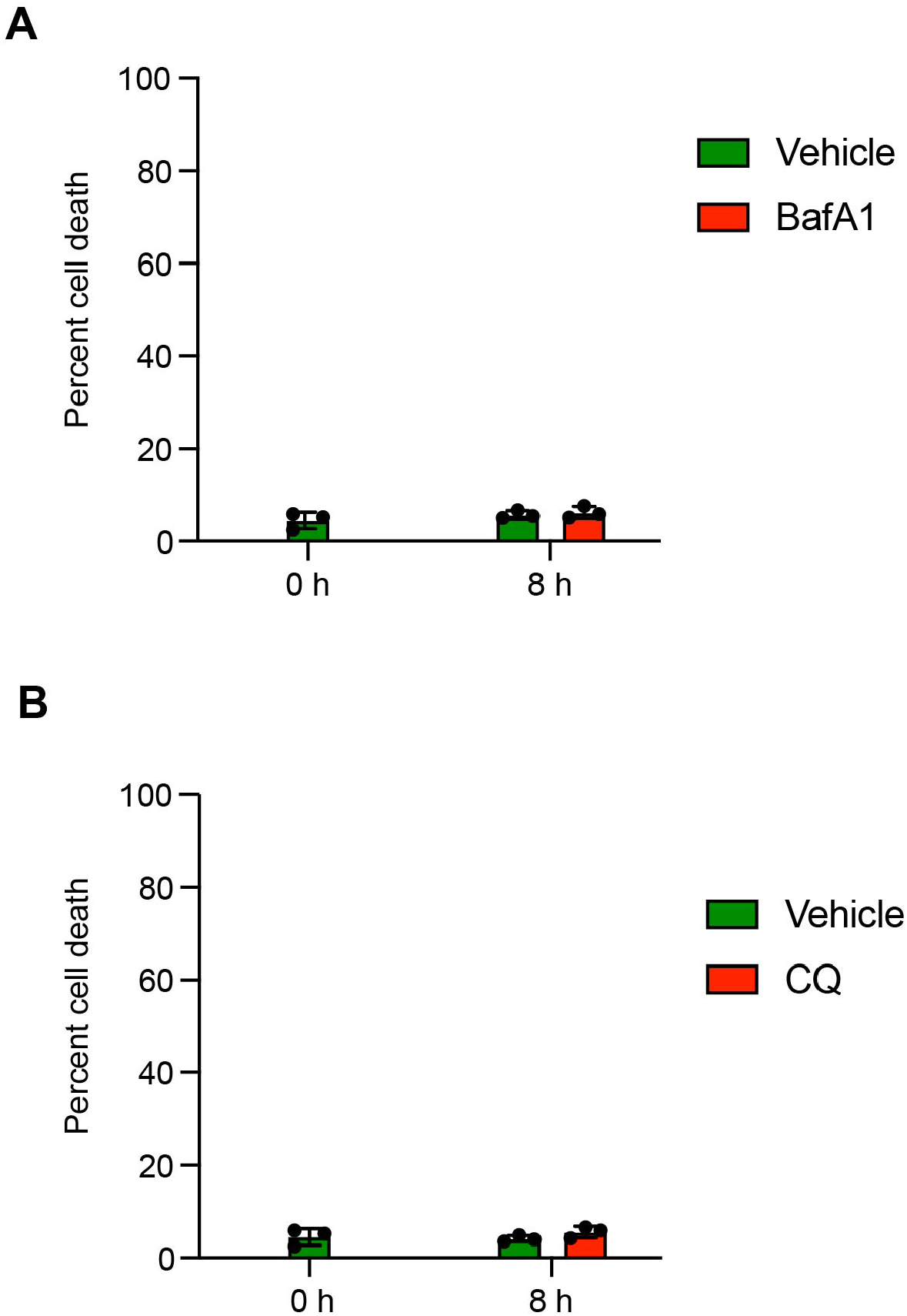
Lysosomal inhibition has a negligible impact on cell death. **(A)** Quantification of cell death in wild-type cells prior to treatment (0 h) or treated with vehicle in serum-free media (Vehicle), or 20 nM BafA1 in serum-free media (BafA1) for 8 h using trypan blue staining (mean ± s.e.m.; n=3). **(B)** Quantification of cell death in wild-type cells prior to treatment (0 h) or treated with vehicle in serum-free media (Vehicle), or 25 µM chloroquine (CQ) in serum-free media (BafA1) for 8 h using trypan blue staining (mean ± s.e.m.; n=3).

**Supplemental Figure 3.**
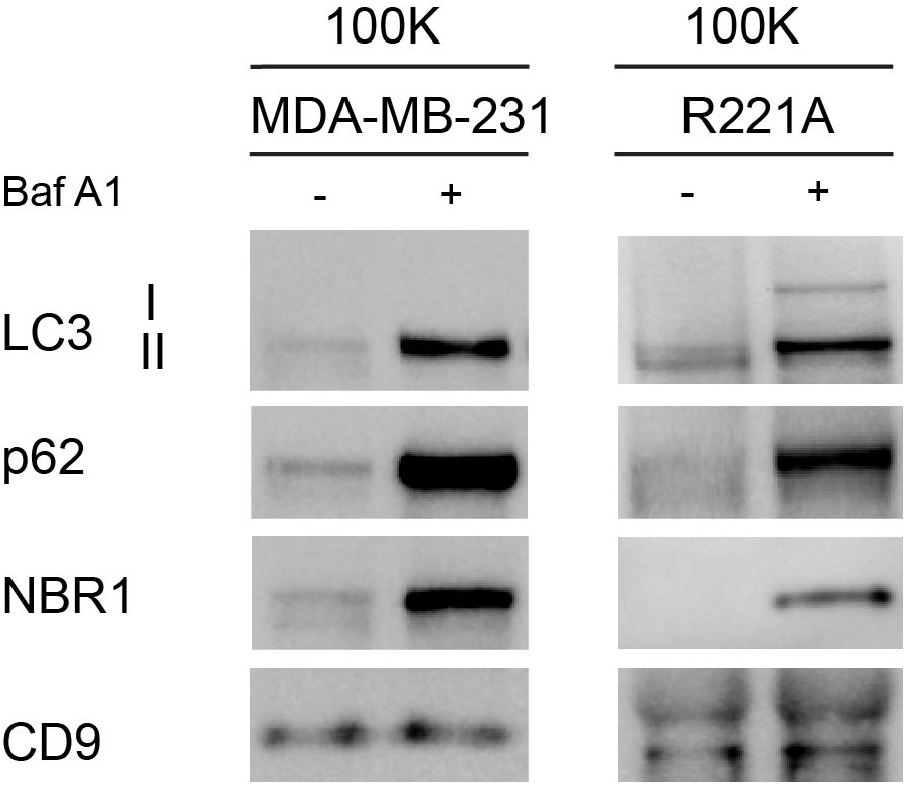
LC3 and autophagy cargo receptors are secreted in EVPs from diverse cell-types in response to lysosome inhibition. 100,000g EVP fractions (100K) from wild-type human MDA-MB-231 and murine R221 breast cancer cell lines treated with vehicle or 20 nM BafA1 in serum-free media for 16 h were collected and immunoblotted for indicated proteins (n=3).

**Supplemental Figure 4.**
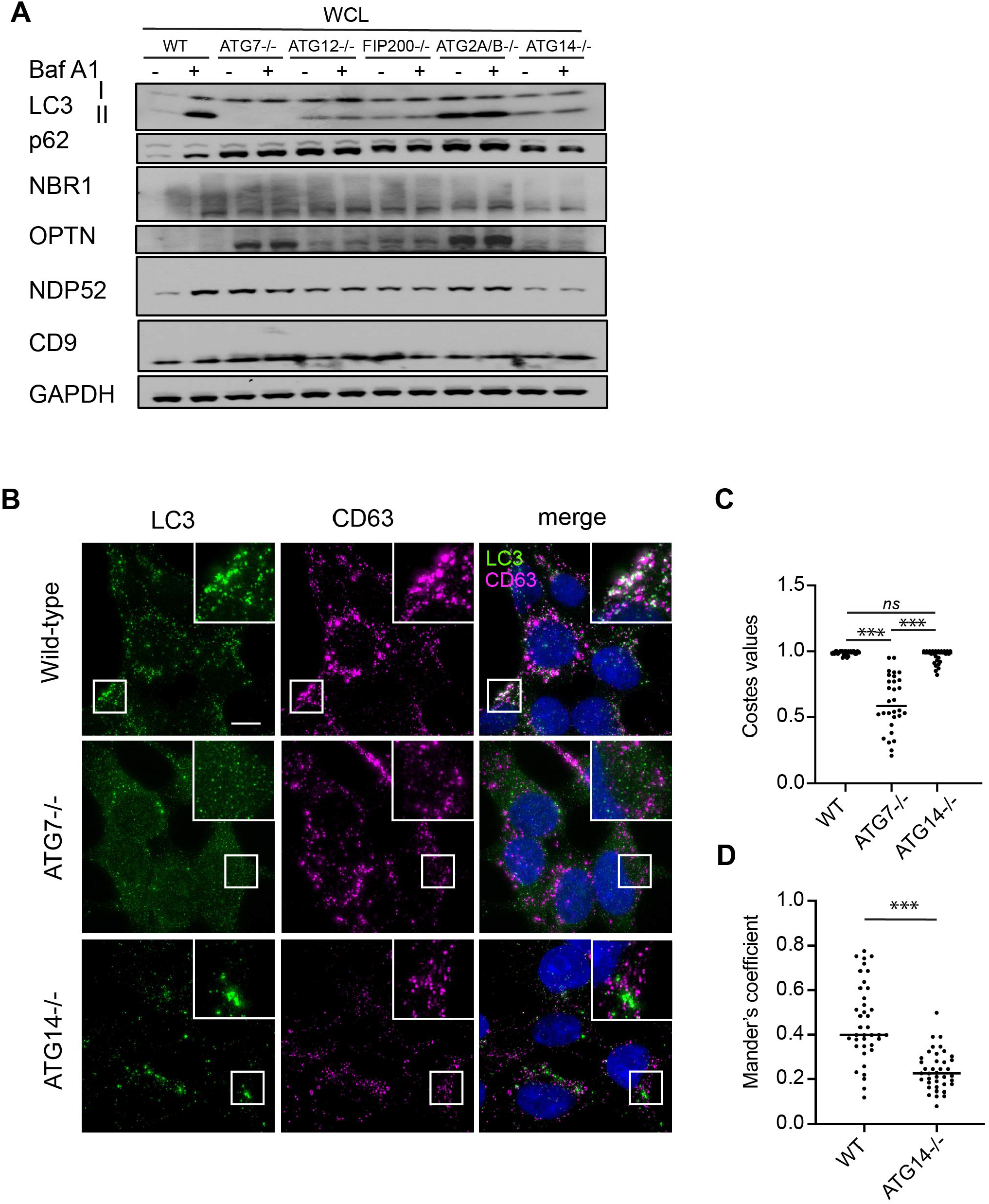
Autophagosome formation is required for autophagy cargo receptor secretion in response to lysosome inhibition. **(A)** Whole cell lysate (WCL) from serum starved HEK293Ts of the indicated genotypes treated with or without 20 nM BafA1 and corresponding to secretion experiments in Figure 5A,B were collected and immunoblotted for LC3, autophagy cargo receptors, CD9 and GAPDH. **(B)** Representative fluorescence micrographs from serum starved wild-type, ATG7−/− and ATG14−/− cells treated with 20 nM BafA1 and immunostained for endogenous LC3 (green) and CD63 (magenta). Scale bar, 10 µm. **(C)** A scatter plot of the P values obtained from Costes significance tests to assess whether the overlap of LC3 and CD63 staining observed in C exceeds thresholds of random co-occurrence. Statistical significance was calculated by one-way ANOVA with Tukey’s post hoc test (mean ± s.e.m.; WT, n = 30; ATG7−/−, n = 30; ATG14−/−, n = 30; ns=not significant, p<0.005=***). **(D)** A scatter plot of Mander’s coefficients for the co-occurrence of LC3 with CD63 in BafA1 treated wild-type and ATG14-/- cells in C. Statistical significance was calculated by an unpaired two-tailed t-test (mean ± s.e.m.; WT, n = 39; ATG14−/−, n = 38; p<0.005=***).

**Supplemental Figure 5.**
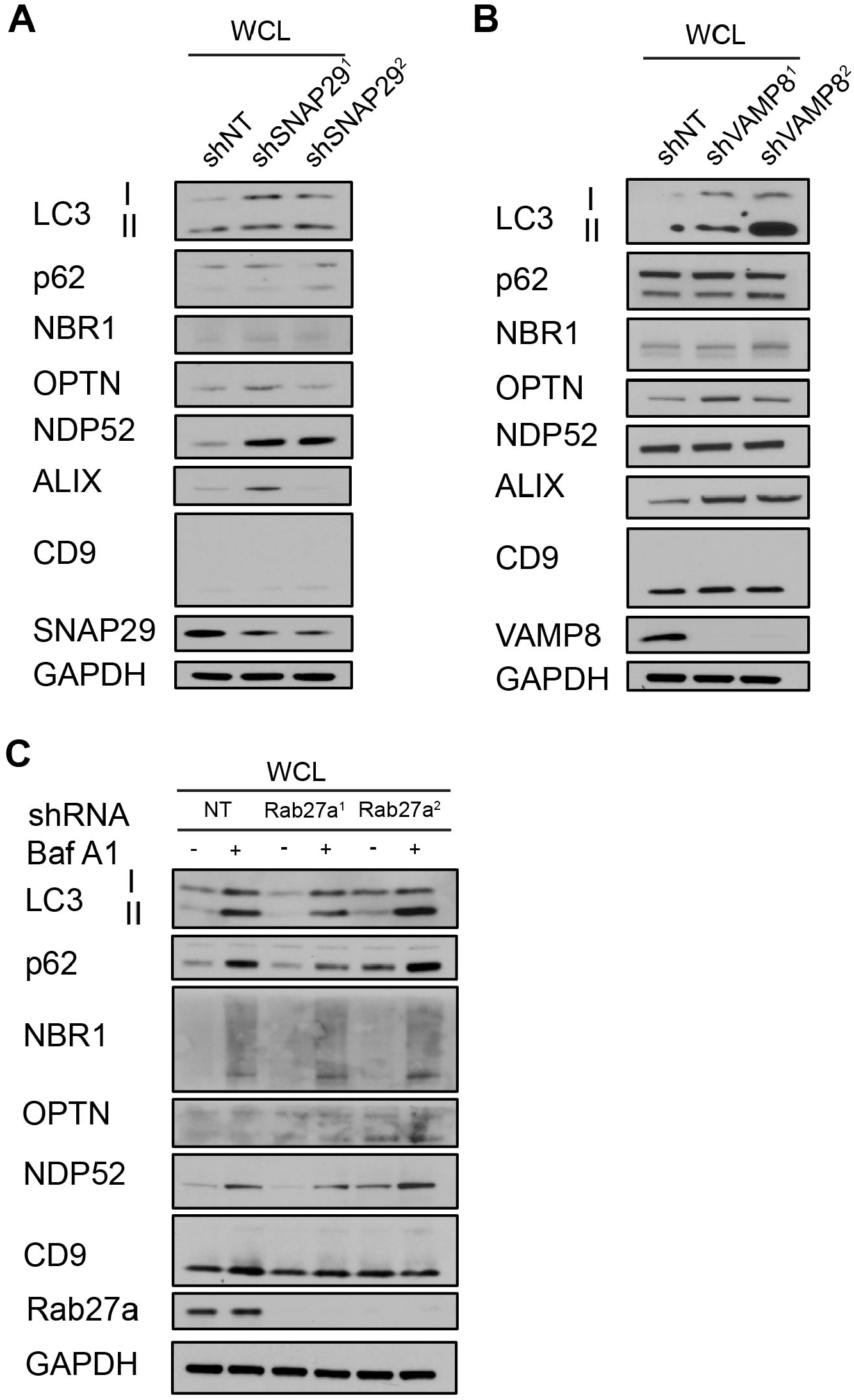
Rab27a is required for autophagy cargo receptor secretion in response to lysosome inhibition. **(A)** WCL from serum starved HEK293Ts that stably express shRNAs targeting SNAP29 (shSNAP29^1^; shSNAP29^2^) or control shRNA (shNT) and corresponding to secretion experiments in Figure 6E,F were immunoblotted for the indicated proteins. **(B)** WCL from serum starved HEK293Ts that stably express shRNAs targeting SNAP29 (shSNAP29^1^; shSNAP29^2^) or control shRNA (shNT) and corresponding to secretion experiments in Figure 6E,F were immunoblotted for the indicated proteins. **(C)** WCL from serum starved HEK293Ts that stably express shRNAs targeting Rab27a (Rab27a^1^; Rab27a^2^) or control shRNA (NT) and corresponding to secretion experiments in Figure 6E,F were immunoblotted for the indicated proteins.

## Notes

**Potential conflicts of interest:** JD is member of the Scientific Advisory Board of Vescor Therapeutics, LLC.

